# Low intra-microbiota antagonism underlies stable and protective barley root microbiota

**DOI:** 10.64898/2026.05.23.727129

**Authors:** Lisa Mahdi, Natalie Peltz, Sinaeda Anderssen, Stanislav P. Kopriva, George Alskief, Gregor Langen, Alga Zuccaro

## Abstract

Plants rely on root-associated microbiota for growth and pathogen protection, yet the community properties underlying stable beneficial functions remain unclear. Here, we show that low intra-microbiota antagonism is associated with stability and pathogen protection in the barley root microbiota. Using a comprehensive culture collection of barley root endophytes, we reconstructed bacterial synthetic communities (SynComs) and compared a core SynCom resembling the natural microbiota with a trait-prioritized SynCom assembled from strains selected for predicted host benefits. Although strains from both communities exhibited broad pathogen-inhibitory potential, the communities behaved differently in planta. The core SynCom showed low internal antagonism, remained stable, and protected barley roots against fungal pathogens without impairing growth. In contrast, the trait-prioritized SynCom exhibited strong internal antagonism, unstable assembly, loss of protection, and reduced root growth. Together, our findings indicate that microbiota function cannot be predicted from individual strain traits alone. Rather, low intra-microbiota antagonism promotes community stability and enables protective functions to emerge in barley roots.

**Significance statement:** Plants rely on root microbiota for disease protection, yet the principles governing community stability and protective function remain poorly understood. Here, we show that low intra-microbiota antagonism is associated with a stable and protective barley root microbiota, whereas strong internal antagonism destabilizes communities, impairs root growth, and abolishes pathogen protection. These findings show that microbiota function depends not only on beneficial traits of individual strains, but also on compatibility among community members.

**Highlights:**

- Low intra-microbiota antagonism is associated with stable and protective barley root microbiota.
- Trait-prioritized SynCom assembly with higher intra-microbiota antagonism can lead to dysbiosis.
- A highly antagonistic *Pseudomonas* strain is buffered by the core SynCom but detrimental in an unstable community.
- Community compatibility better explains protection than individual strain traits.
- Stable bacterial communities support fungal endophyte-mediated protection and maintain pathogen protection across barley genotypes.

## Introduction

Plants live in close association with complex microbial communities, collectively termed the plant microbiota. These communities contribute to plant growth and health by supporting nutrient acquisition, modulating hormone balance, and increasing resistance to biotic and abiotic stress. In this way, the microbiota can function as an extension of the plant immune system ^1–3^. At the same time, plants actively shape their associated microbiota through root exudates, immune signaling, and selective recruitment mechanisms, making the host an important driver of microbiome composition and homeostasis ^3–7^. This reciprocal relationship has led to the concept of the plant and its microbiota as an integrated ecological unit, the holobiont ^8,9^.

The holobiont framework emphasizes that plant health is shaped not only by individual microbial partners, but also by their collective behavior. Many plant-beneficial functions do not arise from single microbes in isolation, but emerge from interactions within the community. Such emergent properties depend on community composition, functional redundancy, and synergistic activities among microbial members ^10–12^. Stable and protective microbiota are therefore thought to rely on compatible microbial interactions that allow community members to coexist while maintaining beneficial functions for the host ^13,14^. However, the ecological principles that define stable and protective microbiota remain poorly understood because community assembly and function depend on dynamic, context-dependent interactions among microbes and between microbes and the host ^15–19^.

When microbiota homeostasis is disrupted, plant-associated communities can shift toward dysbiotic states. Environmental perturbations such as climate change, intensive agricultural practices, monocultures, and the overuse of agrochemicals can alter microbial community structure and reduce beneficial functions ^20–25^. Dysbiotic microbiota are increasingly recognized as associated with disease susceptibility, yield decline, reduced diversity, loss of protective functions, and dominance of opportunistic or pathogenic taxa ^20,26–29^. Despite the importance of this contrast, the community-level interaction patterns that distinguish stable, protective microbiota from dysbiotic states remain largely unresolved ^30^.

Synthetic microbial communities provide a powerful approach to address this knowledge gap. By reconstructing microbial communities under defined conditions, SynComs make it possible to test how plant–microbe and microbe–microbe interactions shape community stability and host protection. Most SynCom studies have focused on identifying beneficial strains or functional traits. However, less is known about how interactions among community members influence whether predicted beneficial functions are maintained, lost, or altered *in planta*.

Here, we used barley as a model to test how intra-microbiota antagonism influences microbiota stability and pathogen protection. We established a comprehensive culture collection of root-associated bacteria from the cultivated barley genotype Golden Promise (GP) and the wild accession HID4 (*Hordeum vulgare* ssp. *spontaneum*) grown under identical soil conditions. From this resource, we assembled two bacterial SynComs: a core SynCom resembling the natural microbiota of cultivated barley and a trait-prioritized SynCom composed of strains selected for predicted plant-beneficial traits. This design allowed us to compare a community reflecting natural coexistence with one assembled based on expected functional potential. We show that beneficial traits of individual bacterial strains alone do not predict community-level function. Instead, low antagonism among microbiota members is associated with community stability, limits disruptive competition, and enables protective functions to emerge in barley roots.

## Results

### Barley host genotype contributes to root microbiota composition

To reconstruct barley root microbiota under defined conditions, we first characterized the bacterial communities associated with two genetically distinct barley accessions: the cultivated genotype GP and the wild accession HID4. Because plant-associated microbiota are shaped by both environmental and host-genetic factors, including domestication and breeding history ^31–33^, we grew both accessions under identical soil conditions and profiled their root-associated bacterial communities. In parallel, we established a culture collection of bacterial root endophytes from the same samples, providing the experimental resource for subsequent SynCom assembly and manipulation experiments (Fig. 1A).

**Figure 1:**
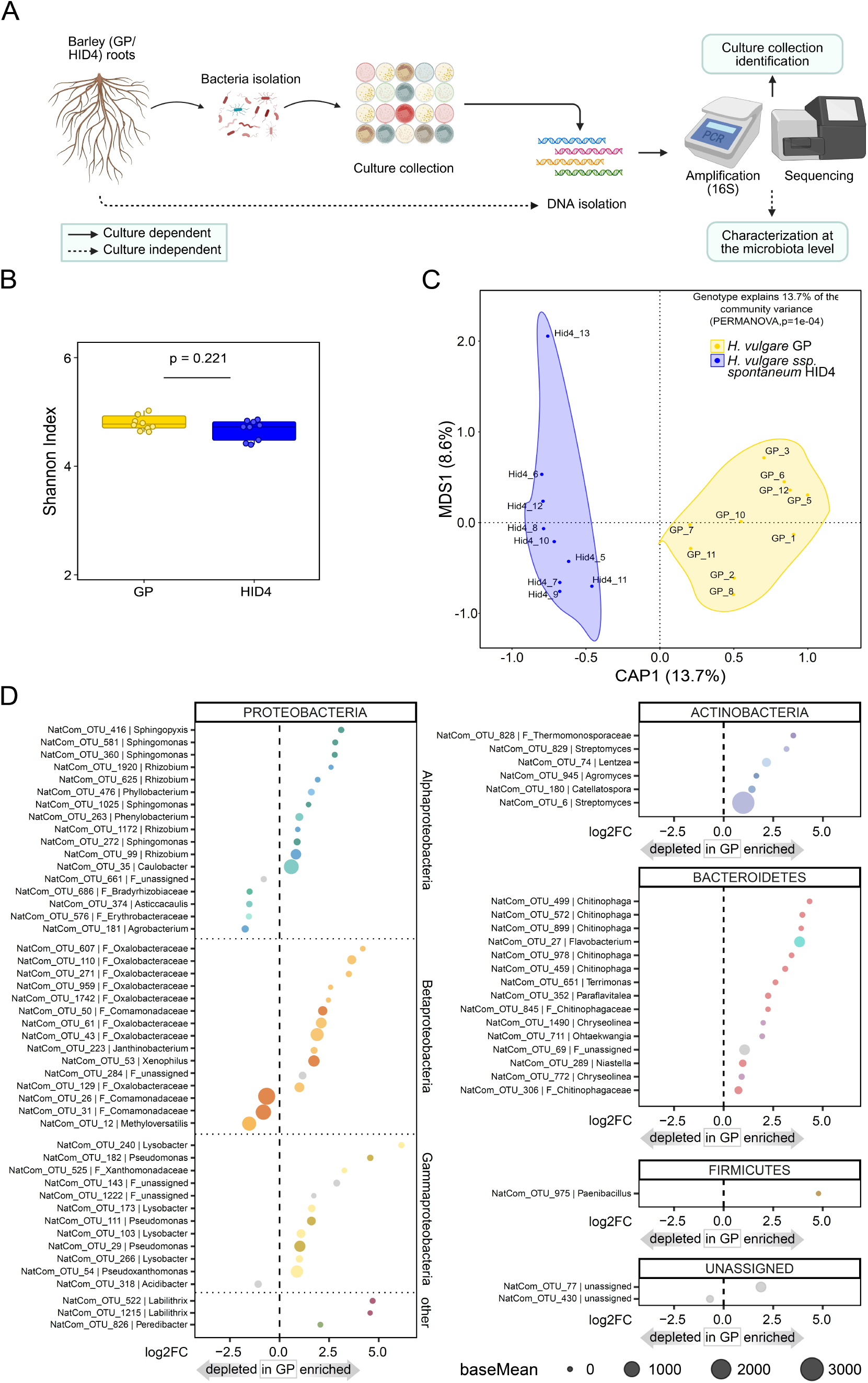
Root bacterial microbiota of barley accessions GP and HID4 differ in community composition and taxon-specific abundance. **A)** Schematic overview of the culture-dependent and culture-independent approaches used to characterize barley root-associated bacterial communities and establish the culture collection. **B)** Shannon diversity index of the root bacterial microbiota associated with GP (yellow) and HID4 (blue). Differences between genotypes were tested using a non-parametric Mann–Whitney U test. **C)** Constrained principal component analysis of bacterial community composition in GP and HID4 roots. Differences in community composition were assessed using PERMANOVA. **D)** Log₂ old changes of bacterial OTUs significantly enriched or depleted in GP roots compared with HID4 roots, grouped by phylum. Differentially abundant OTUs were identified after filtering out OTUs absent from more than 90% of samples, followed by DESeq2 analysis. Only OTUs showing significant enrichment or depletion in GP relative to HID4 are displayed. Dot size represents the DESeq2 baseMean, corresponding to the mean normalized abundance across all samples. Dot color indicates taxonomic assignment at the family level.

The resulting culture collection comprised 1,926 bacterial strains representing 313 OTUs (Table S1). To compare the natural root microbiota of GP and HID4, we generated bacterial community profiles from individual barley roots by amplicon sequencing of the V5–V7 region of the 16S rRNA gene. Alpha-diversity did not differ significantly between GP and HID4, consistent with previous studies comparing cultivated and wild barley genotypes (Fig. 1B, Fig. S1A) ^32^. In contrast, host genotype had a small but significant effect on bacterial community composition, explaining 13.7% of the variation in beta-diversity (Fig. 1C; PERMANOVA, p = 1 × 10⁻⁴; Fig. S1B). This difference was mainly associated with OTUs affiliated with Proteobacteria, Bacteroidetes, and Actinobacteria (Fig. S2).

Differential abundance analysis identified 73 significantly enriched OTUs between GP and HID4 roots (|log₂ fold change| ≥ 0.584, padj ≤ 0.05), with most enriched OTUs associated with GP roots (63 OTUs) rather than HID4 roots (10 OTUs; Table S2). Several GP-enriched OTUs belonged to bacterial genera commonly associated with endophytic or rhizospheric lifestyles and reported plant-beneficial functions (Fig. 1D). These included proteobacterial genera such as *Sphingomonas*, *Pseudoxanthomonas*, *Rhizobium*, *Caulobacter*, *Lysobacter*, and *Pseudomonas* ^34–41^, as well as actinobacterial genera such as *Streptomyces*, *Lentzea*, and *Catellatospora* ^42,43^. Additional GP-enriched OTUs were affiliated with Chitinophagaceae, a family previously linked to plant growth promotion and suppression of fungal pathogens, including *Rhizoctonia solani* and *Bipolaris sorokiniana* (*Bs*) ^44–48^. Together, these results show that barley host genotype contributes to root microbiota composition under the same soil conditions.

From this resource, we identified core taxa in both accessions that were consistently detected in every analyzed plant, comprising 235 core taxa in GP and 239 core taxa in HID4 (Tables S3, S4). The identification of core taxa as well as bacterial groups previously linked to plant-beneficial functions provided the basis for assembling representative bacterial SynComs to test how community structure influences stability and protection.

### Intra-microbiota antagonism distinguishes stable from dysbiotic synthetic communities

Synthetic microbial communities provide a controlled framework to link microbiota composition with community function. To distinguish the contribution of individual strain traits from community structure, we assembled two bacterial SynComs from the barley root culture collection. *Hv*SC1 was designed to resemble the natural core microbiota of the cultivated barley genotype GP. This community was assembled from cultured isolates corresponding to bacterial OTUs consistently detected across all GP root samples grown in Cologne soil (n = 10), thereby representing prevalent members of the native GP root microbiota (Fig. 2A, B). From this core set, we selected isolates that collectively captured the taxonomic composition of the natural GP root microbiome. To assess whether these taxa are broadly associated with barley, we also analyzed root microbiota from six additional barley species, including multiple accessions and both wild and domesticated lines (Table S5). *Hv*SC1 taxa were detected across multiple accessions, indicating that they represent frequent members of the barley-associated microbiota (Fig. 2C).

**Figure 2:**
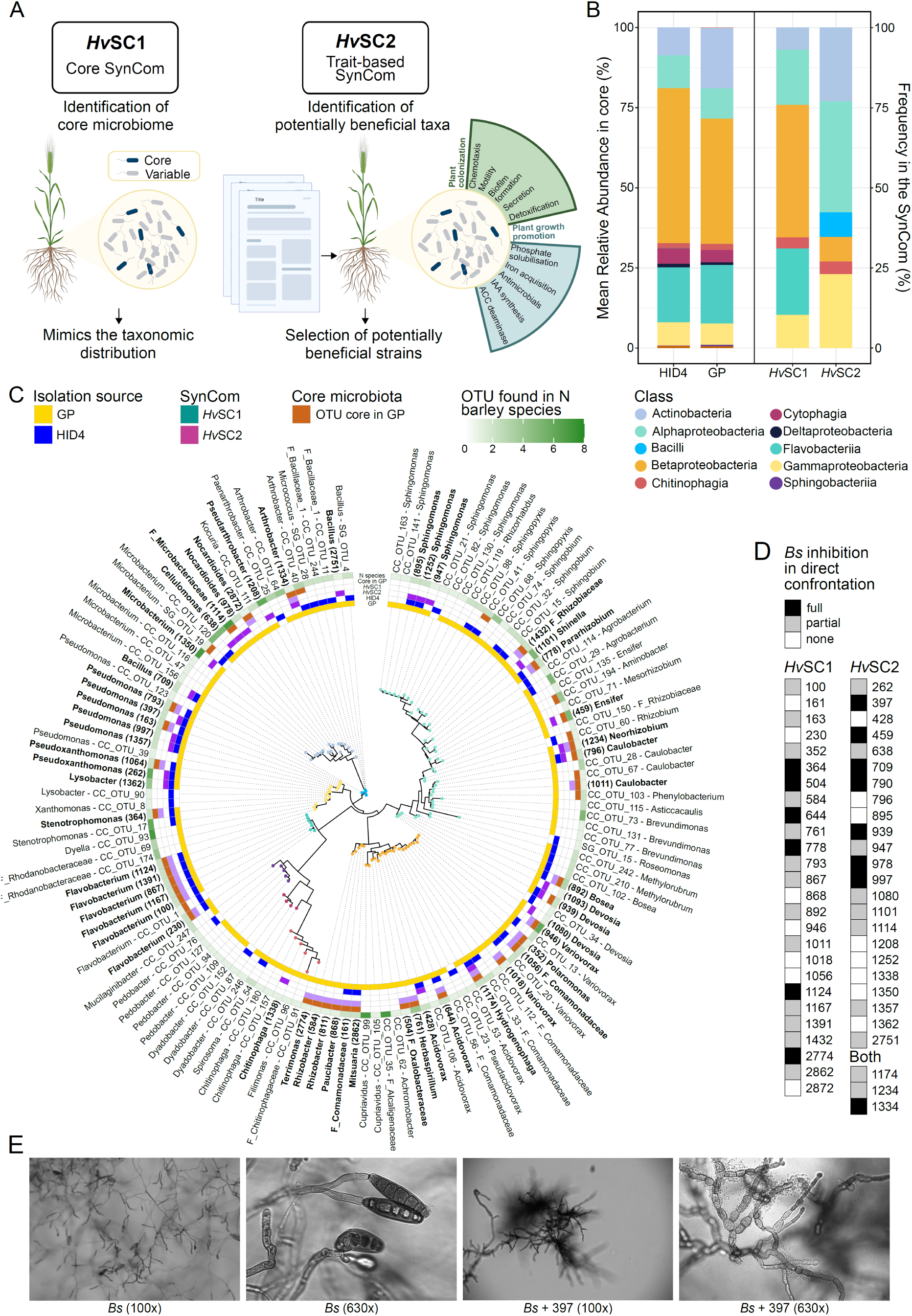
Core microbiota reconstruction and trait-prioritized strain selection generate taxonomically distinct barley SynComs with broad pathogen-inhibitory potential. **A)** Schematic overview of the two strategies used to assemble bacterial synthetic communities. *Hv*SC1 was designed as a core SynCom resembling the natural GP root microbiota by selecting cultured isolates corresponding to core OTUs and matching the taxonomic distribution of the native community. *Hv*SC2 was designed as a trait-prioritized SynCom by selecting GP-derived isolates affiliated with bacterial groups previously associated with plant-beneficial functions. **B)** Class-level taxonomic composition of natural root microbiota and assembled SynComs. The left panel shows the relative abundance of bacterial classes in the natural root microbiota of HID4 and GP. The right panel shows the class-level composition of *Hv*SC1 and *Hv*SC2. Each OTU is represented with equal weight in the SynCom bars. **C)** Phylogenetic relationships among culture collection isolates including all members of *Hv*SC1 and *Hv*SC2. Only pure culture collection members are included. Rings indicate isolation source, SynCom membership, GP core microbiota status as inferred though amplicon sequencing, and the number of barley species in which each OTU was detected. Tree-tip labels indicate genus-level taxonomy when available; otherwise, family-level taxonomy is shown using the prefix “F_”. SynCom strains are annotated with their corresponding strain identifiers in bold. Non-members include the corresponding OTU ID. **D)** Pathogen-inhibitory capacity of individual *Hv*SC1 and *Hv*SC2 strains against *Bipolaris sorokiniana* (*Bs*) in direct confrontation assays. Strains are ranked by inhibition strength. Colors indicate full inhibition, partial inhibition, or no inhibition. **E)** Microscopy images of *Bs* grown alone or in confrontation with *Pseudomonas* strain 397. Images show *Bs* hyphae and bacterial accumulation near fungal hyphae at 100× and 630× magnification.

*Hv*SC2 was assembled using a trait-prioritized strategy. It comprised GP-derived bacterial isolates selected based on taxonomic affiliation with genera previously associated with plant growth promotion or pathogen suppression. Thus, *Hv*SC2 emphasized predicted strain-level beneficial potential rather than prevalence, natural co-occurrence, or representation of native microbiota structure (Fig. 2A, B). In contrast to *Hv*SC1, only seven *Hv*SC2 strains corresponded to OTUs defined as core in GP or in the additionally tested barley accessions (Fig. 2C).

We next determined the pathogen-inhibitory potential of each SynCom strain in direct confrontation assays against the fungal pathogen *Bs*. Pathogen inhibition was common among barley root-associated bacteria in both SynComs, despite their different assembly strategies. In total, 75% of *Hv*SC1 strains (22/29) and 69% of *Hv*SC2 strains (18/26) inhibited *Bs* growth (Fig. 2D). Strong inhibition was particularly evident for strains such as *Pseudomonas* strain 397, where *Bs* suppression was visually pronounced and accompanied by dense bacterial accumulation near fungal hyphae (Fig. 2E). Notably, *Bs* inhibitory potential could not be inferred from the prevalence or genomic copy number of specific biosynthetic gene clusters (BGCs) (Fig. S3A, B).

We then investigated the effects of both SynComs on their original host, the barley cultivar GP. At 6 days post inoculation (dpi), inoculation with *Hv*SC1 did not affect root biomass. In contrast, *Hv*SC2 significantly reduced root fresh weight, despite being assembled from strains with predicted plant-beneficial traits (Fig. 3A).

**Figure 3:**
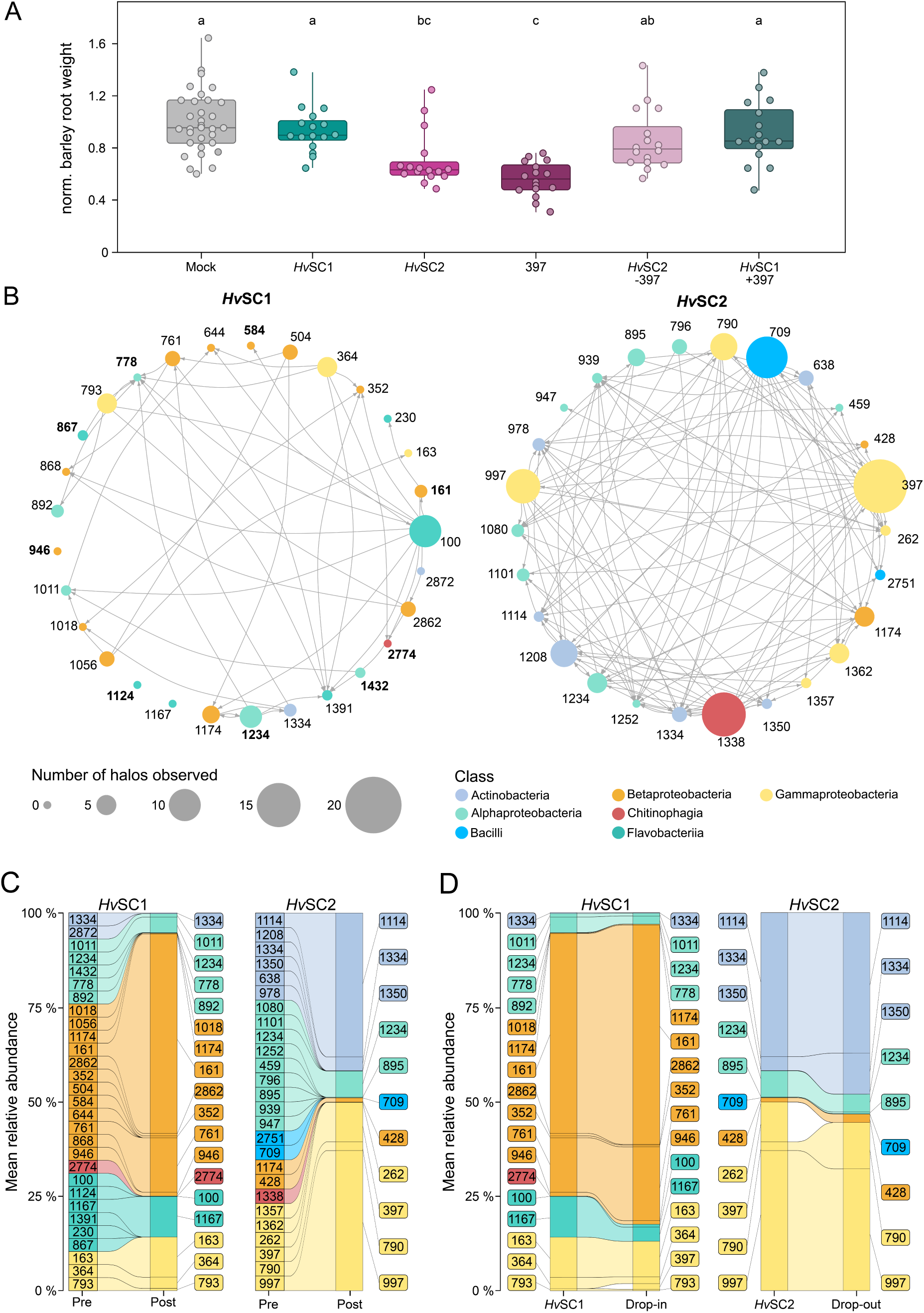
Intra-SynCom antagonism shapes community stability, antagonist dominance, and barley root growth. **A)** Root fresh weight of barley cultivar GP inoculated with *Hv*SC1, *Hv*SC2, *Pseudomonas* strain 397, and bacterial mixtures from drop-out and drop-in experiments. Statistical differences among treatments were assessed using a Kruskal–Wallis test followed by Dunn’s multiple-comparison test (p < 0.05). **B)** Bacterial inhibition networks of *Hv*SC1 and *Hv*SC2. Arrows indicate inhibitory interactions and point from the inhibiting strain to the inhibited strain. Node colors indicate bacterial classes according to the legend, and node size reflects the number of inhibition halos produced by each strain. **C)** Mean relative abundance of *Hv*SC1 and *Hv*SC2 strains in barley GP roots at inoculation and at the experimental endpoint, 6 days post inoculation. Colors of strata and alluvial flows indicate bacterial classes as shown in the legend. **D)** Mean relative abundance of *Hv*SC1 and *Hv*SC2 strains in barley GP roots at 6 days post inoculation in the original SynComs and corresponding drop-in or drop-out treatments with *Pseudomonas* strain 397. Colors indicate bacterial class. Relative abundances in **C** and **D** were first computed per sample and then averaged across treatment groups. The per-sample relative abundances are shown in Fig. S6.

We next tested whether this functional difference was associated with community interaction structure and stability. Using Burkholder plate assays, we quantified inhibitory interactions among SynCom members across 2,260 pairwise combinations. Antagonism was widespread in *Hv*SC2 but much less frequent in *Hv*SC1. *Hv*SC2 showed 115 inhibitory interactions, corresponding to an average of 4.4 inhibitions per strain, whereas *Hv*SC1 showed 52 inhibitory interactions, corresponding to 1.8 inhibitions per strain (Fig. 3B). Thus, the trait-prioritized SynCom displayed substantially stronger internal antagonism than the core SynCom.

To test whether differences in intra-community antagonism were associated with community stability in planta, we performed amplicon sequencing of inoculated GP roots. *Hv*SC1 remained more stable than *Hv*SC2: a larger proportion of *Hv*SC1 strains persisted in roots (65%) compared with *Hv*SC2 strains (42%), and *Hv*SC1 largely retained its initial class-level taxonomic composition. In contrast, *Hv*SC2 underwent pronounced compositional changes, including loss of Alphaproteobacteria, enrichment of Betaproteobacteria, and dominance by two highly antagonistic *Pseudomonas* strains, 997 and 397 (Fig. 3C).

Together, these results show that strong intra-microbiota antagonism is associated with instability, competitive dominance, and impaired host performance, whereas low antagonism is associated with stable community assembly in barley roots.

### A dominant *Pseudomonas* antagonist impairs host growth but is buffered by the core SynCom

The instability of *Hv*SC2 was accompanied by the dominance of two highly antagonistic *Pseudomonas* strains, including strain 397 (Fig. 3B, C). Because strain 397 reached high relative abundance in *Hv*SC2 and showed strong inhibitory activity against other SynCom members, we next tested whether it contributed to the reduced host growth caused by the trait-prioritized community. When applied alone to the barley cultivar GP, strain 397 largely recapitulated the root growth reduction observed after *Hv*SC2 inoculation (Fig. 3A). It was also the only strain across both SynComs that negatively affected both barley root and shoot weight in mono-association experiments (Fig. S4, S5). These results indicate that strain 397 is sufficient to impair host performance when acting outside a stabilizing community context.

We next tested the effect of strain 397 within each SynCom. Removing strain 397 from *Hv*SC2 in a drop-out experiment mitigated the negative effect of this community on root biomass (Fig. 3A), indicating that strain 397 contributes substantially to the deleterious host phenotype of *Hv*SC2. In contrast, adding strain 397 to *Hv*SC1 did not impair root growth (Fig. 3A). Thus, the same antagonistic strain had different effects depending on community context. Consistent with this buffering effect, strain 397 reached lower relative abundance when introduced into *Hv*SC1 (1.5%) than when present in *Hv*SC2 (10.5%) (Fig. 3D). This suggests that *Hv*SC1 constrains the proliferation of strain 397 *in planta*. Notably, strain 397 inhibited several *Hv*SC1 members in pairwise Burkholder plate assays, whereas none of the *Hv*SC1 strains individually inhibited strain 397 (Fig. 3B). Therefore, containment of strain 397 in *Hv*SC1 is unlikely to result from a single direct antagonistic interaction. Instead, it appears to emerge from the collective community context.

Together, these results show that a dominant antagonistic strain can impair host growth in an unstable, highly antagonistic community but can be buffered within a stable core SynCom. This supports the idea that community compatibility limits competitive dominance and preserves host-compatible microbiota function.

### Genomic biosynthetic potential alone does not explain intra-microbiota antagonism

Microbial antagonism is often mediated by specialized secondary metabolites encoded within biosynthetic gene clusters (BGCs) ^49,50^. Thus, we investigated whether the differences in antagonism between *Hv*SC1 and *Hv*SC2 could be explained by genomic biosynthetic potential. To address this, we sequenced the genomes of SynCom strains and examined the relationships among BGC repertoire, phylogenetic distance, and bacterial inhibition profiles. In Streptomycetaceae, inhibition phenotypes can be predicted by BGC similarity and phylogenetic distance ^51^. We therefore tested whether similar relationships were present across our phylogenetically diverse barley root bacterial collection.

Using BiG-SCAPE, we grouped predicted BGCs from SynCom strain genome assemblies into gene cluster families (GCFs, Fig. S7 ^52^). Phylogenetic distance was a strong predictor of GCF dissimilarity across the strain collection (Fig. S7A), consistent with evolutionary divergence in secondary metabolite potential ^53^. However, neither phylogenetic distance nor GCF similarity predicted inhibition profile dissimilarity in *Hv*SC1 or *Hv*SC2 (Fig. S7B-E). These relationships remained non-significant after accounting for the correlation between phylogeny and BGC profile (Table S6). Together, these results indicate that the stronger antagonism observed in *Hv*SC2 cannot be explained by phylogenetic relatedness among strains.

We then asked whether antagonistic output within each SynCom was associated with strain-level BGC repertoire. Total BGC count correlated positively with the number of inhibition halos produced by *Hv*SC2 strains (r = 0.77, p < 0.001; Fig. 4B), but not by *Hv*SC1 strains (r = 0.12, p = 0.546; Fig. 4A). Thus, in the more antagonistic *Hv*SC2 community, strains with larger biosynthetic repertoires tended to display stronger inhibitory activity, whereas this relationship was absent in the core SynCom.

**Figure 4:**
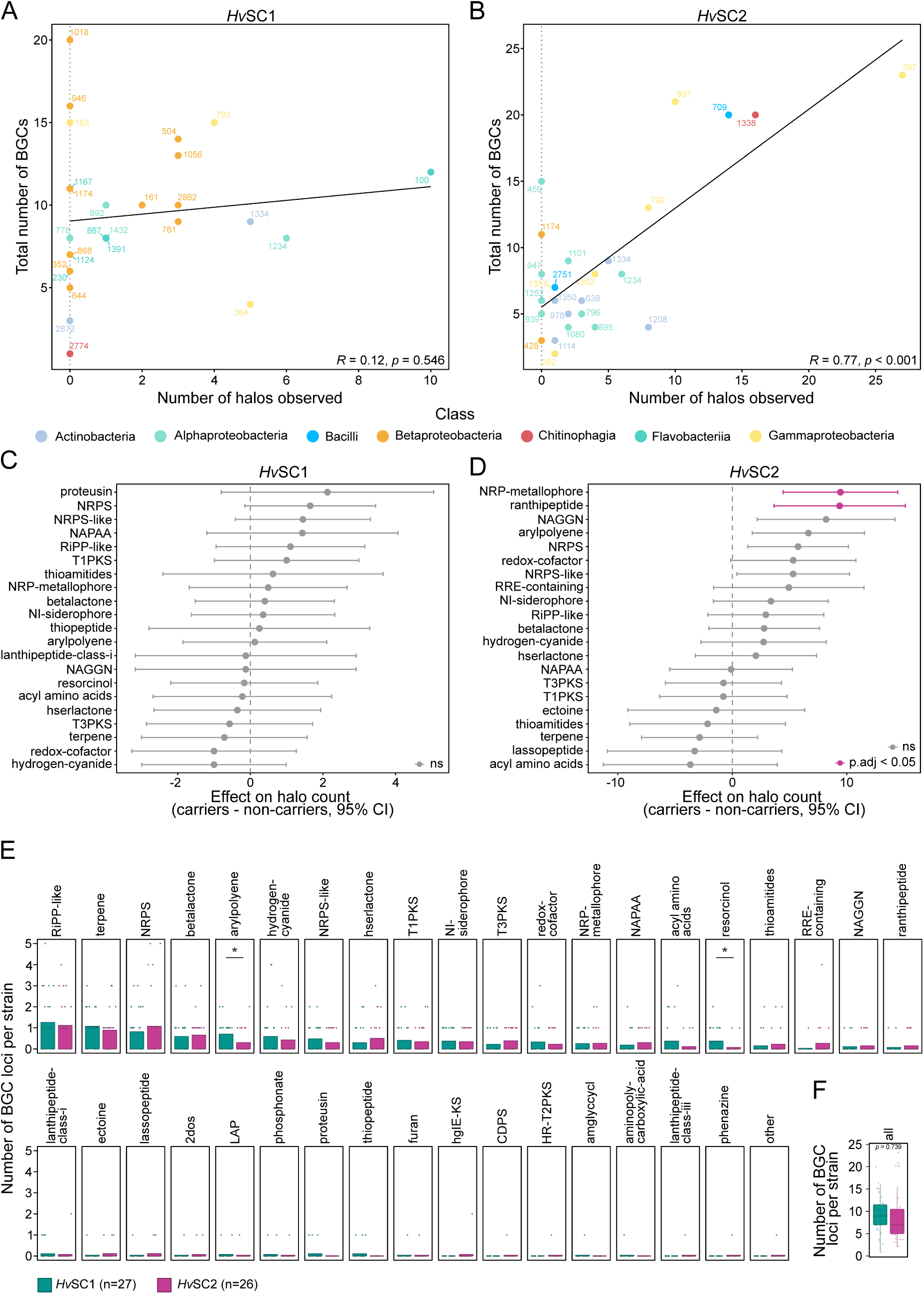
Biosynthetic potential predicts antagonism in *Hv*SC2 but does not explain SynCom-wide differences in inhibition. A–B) Relationship between total BGC count per strain and the number of inhibitory halos produced by *Hv*SC1 (**A**) and *Hv*SC2 (**B**) strains. Each dot represents one strain and is colored by bacterial class. Linear regressions are shown, with Pearson correlation coefficients and associated p-values indicated. **C–D)** Estimated effects of individual BGC types on halo production in *Hv*SC1 (**C**) and *Hv*SC2 (**D**), derived from separate linear models. The x-axis shows the estimated difference in mean halo count between strains carrying or lacking each BGC type. Error bars indicate 95% confidence intervals. BGC types with BH-adjusted p < 0.05 are highlighted in pink. **E)** Distribution of per-strain BGC locus counts across BGC types in *Hv*SC1 and *Hv*SC2. Each facet represents one BGC type, ordered by total locus count. Bars show group means, and dots represent individual strains. Asterisks indicate significant differences between SynComs based on Welch’s two-sample t-test without multiple-testing correction (*p < 0.05, **p < 0.01, ***p < 0.005). **F)** Total number of BGC loci per strain compared between *Hv*SC1 and *Hv*SC2 strains. Statistically significant differences were tested using a Welch’s two-sample t-test.

To identify BGC types associated with antagonistic activity, we tested whether individual BGC classes predicted halo production within each SynCom. In *Hv*SC1, no BGC type was significantly associated with halo counts (Fig. 4C). In *Hv*SC2, however, NRP-metallophore and ranthipeptide clusters were significantly associated with higher halo counts (Fig. 4D). Metallophores can contribute to competition through metal scavenging in resource-limited environments ^54^, whereas ranthipeptides, although mainly studied in intraspecific signaling contexts, may also contribute to microbial interference ^55^. Importantly, these BGC classes were not enriched in *Hv*SC2 relative to *Hv*SC1 (Fig. 4E). Further, overall BGC richness did not differ between the two SynComs (Fig. 4F), suggesting that differences in intra-community antagonism do not result from an enrichment in certain BGC types or greater overall biosynthetic potential alone. The only cluster types that differed in prevalence between the SynComs were arylpolyene and resorcinol clusters, both of which were more abundant in *Hv*SC1 (Fig. 4E). Arylpolyene clusters are primarily associated with oxidative stress protection and biofilm formation ^56^, whereas resorcinol-associated biosynthesis is linked to biocontrol activity against soilborne fungal pathogens ^57^, suggesting a repertoire oriented toward environmental resilience and pathogen suppression rather than intra-community competition.

Together, these findings show that intra-microbiota antagonism is not determined simply by the presence or diversity of BGCs. Rather, the relationship between biosynthetic potential and antagonistic activity appears to be community-dependent, with a strong association in *Hv*SC2 but not in *Hv*SC1. This suggests that ecological interactions may influence whether genomic potential is functionally deployed.

### Microbiota-mediated protection depends on community context rather than strain-level potential

Having shown that the two SynComs differed in internal antagonism, stability, and effects on host growth, we next asked whether these differences also affected their pathogen-protective capacity. We tested the ability of *Hv*SC1 and *Hv*SC2 to protect barley roots against the fungal pathogen *Bs*. To determine whether these effects were specific to the original host genotype, we performed the assays in both the cultivated barley genotype GP and the wild accession HID4. The two SynComs differed strongly in protective capacity. *Hv*SC1 significantly reduced *Bs* colonization in both GP and HID4, indicating that its protective effect was maintained across barley accessions (Fig. 5B). In contrast, *Hv*SC2 failed to reduce *Bs* colonization in either genotype, despite containing many strains with pathogen-inhibitory activity (Fig. 2D, Fig. 5B). Thus, pathogen inhibition by individual strains did not predict protection at the community level.

**Figure 5:**
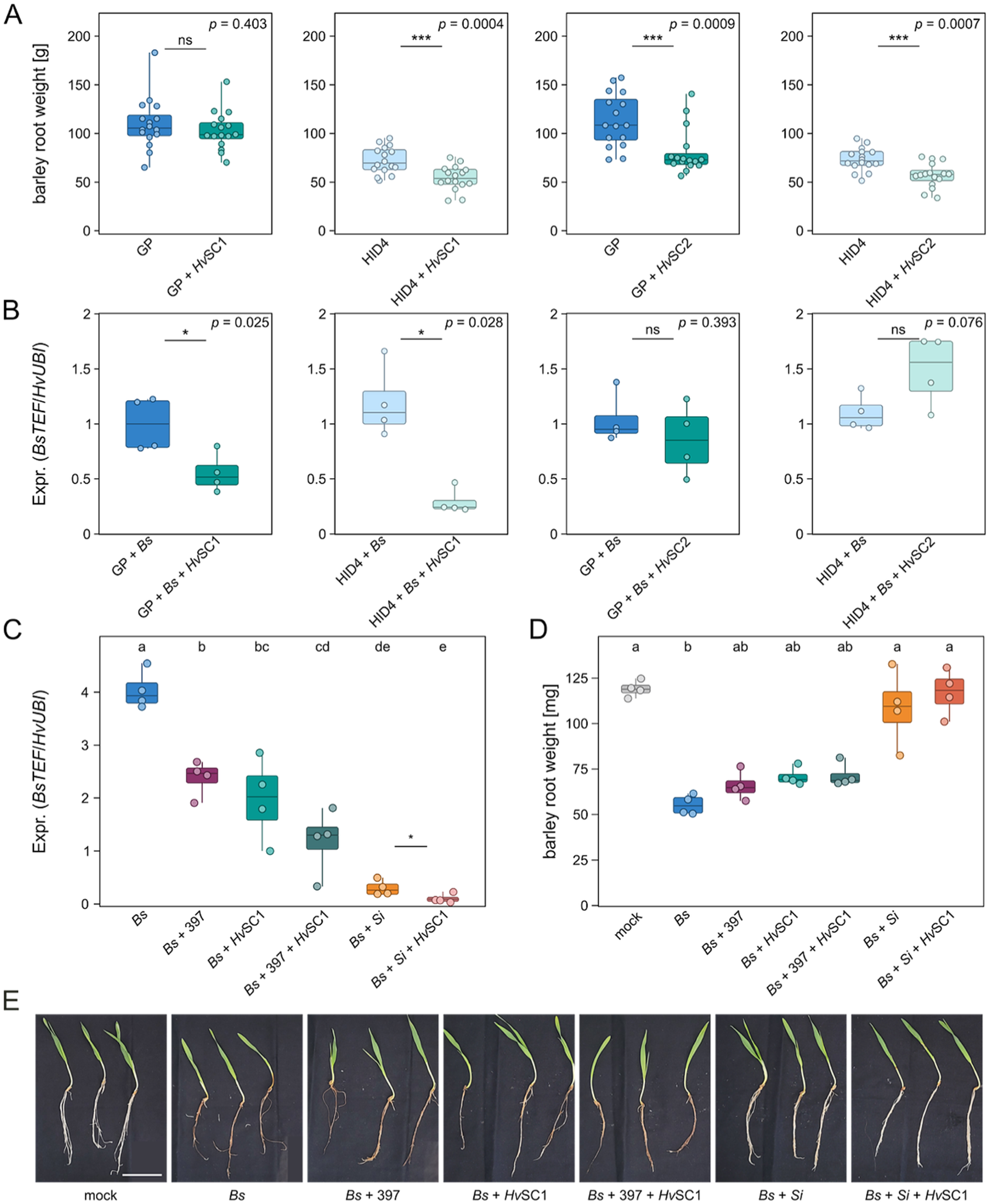
Community context shapes core bacterial, beneficial fungal, and inter-kingdom protection against fungal pathogens. **A)** Root fresh weight of the barley cultivar GP and wild accession HID4 inoculated with *Hv*SC1 or *Hv*SC2. Statistical differences were assessed using two-sample t-tests (p < 0.05). **B)** *Bipolaris sorokiniana* (*Bs*) colonization of GP and HID4 roots inoculated with *Bs* alone or together with *Hv*SC1 or *Hv*SC2, shown relative to *Bs* alone. Statistical differences were assessed using two-sample t-tests (p < 0.05). **C–D)** Relative *Bs* colonization **(C)** and root fresh weight **(D)** of barley cultivar GP inoculated with *Bs* alone or in combination with *Hv*SC1, *Pseudomonas* strain 397, and the beneficial fungal endophyte *Serendipita indica* (*Si*). Statistical differences were assessed using Kruskal–Wallis tests followed by Dunn’s multiple-comparison tests, or ANOVA followed by Tukey’s post-hoc tests, depending on data distribution. Statistical letters indicate significant differences among groups; asterisks indicate pairwise t-tests (*p < 0.05). **E)** Representative images of barley plants inoculated with the indicated microbial treatments at 6 days post inoculation. Scale bar, 5 cm.

The protective capacity of *Hv*SC1 in both barley accessions is consistent with previous reports of largely host-independent microbiota-mediated protection against *Bs* ^58^. Growth responses, however, depended on host genotype. *Hv*SC2 reduced root growth in both GP and HID4, consistent with its dysbiotic behavior. *Hv*SC1 did not affect root growth in its native host GP, but significantly reduced root growth in HID4 (Fig. 5A). These results indicate that microbiota-mediated protection can be maintained across the tested barley accessions, whereas host growth compatibility can be genotype dependent.

We then asked whether the pathogen-inhibitory activity of the dominant antagonistic *Pseudomonas* strain 397 was retained *in planta* and across community contexts. Strain 397 was isolated from barley roots and strongly suppressed *Bs* in direct confrontation assays (Fig. 2E). Moreover, related *Pseudomonas* taxa have previously been associated with biocontrol activity ^59^. When inoculated alone, strain 397 significantly reduced *Bs* colonization of barley roots to levels comparable to *Hv*SC1 (Fig. 5C), showing that strain 397 can confer pathogen protection as an individual strain *in planta*. However, this protective potential does not explain the behavior of the trait-prioritized community. *Hv*SC2 failed to protect against *Bs* even though it contained strain 397 and many other strains with pathogen-inhibitory activity (Fig. 2D, Fig. 5B, C). In contrast, when strain 397 was introduced into *Hv*SC1, pathogen protection was maintained (Fig. 5C, D), indicating that its pathogen-inhibitory capacity is compatible with a stable microbiota context. Together, these results show that protective traits of individual strains can be retained, buffered, or lost depending on community context. Thus, microbiota-mediated protection emerges from community structure rather than from the additive effects of strain-level pathogen inhibition.

### A stable bacterial community enhances protection mediated by a beneficial fungal endophyte

We next asked whether the protective capacity of *Hv*SC1 could extend to an inter-kingdom community context by combining it with the beneficial fungal endophyte *Si*. Consistent with previous reports, *Si* alone strongly reduced colonization by *Bs* ^11,58^ (Fig. 5C, D). This protective effect was further enhanced when *Si* was combined with *Hv*SC1, indicating that the stable bacterial community can support inter-kingdom protection (Fig. 5C, D). Protection mediated by *Hv*SC1, *Si*, or their combination was also reflected at the phenotypic level, including reduced pathogen-induced root weight loss and reduced root browning (Fig. 5D, E).

To determine whether this protection extended beyond the native host genotype and a specific fungal pathogen, we tested the same microbial combinations in HID4 and against *Fusarium graminearum (Fg). Hv*SC1 and *Si* reduced *Bs* colonization in the wild barley accession HID4 and reduced *Fg* colonization, indicating that the protective effect was not restricted to GP or to *Bs* (Fig. S8, S9). Importantly, under simultaneous infection with *Fg* and strain 397, *Hv*SC1 alone reduced *Fg* colonization but failed to rescue host root growth. Phenotypic rescue required the additional presence of *Si*, stressing the relevance of inter-kingdom microbiota in plant protection under strong disease pressure (Fig. S9). Indeed, *Hv*SC1 reduced colonization by both fungal pathogens without impairing colonization by *Si*, suggesting compatibility between core bacterial and beneficial fungal community members within the same root environment (Fig. S10).

Together, these findings show that a stable bacterial SynCom can support beneficial inter-kingdom interactions and enhance host protection. More broadly, they reinforce that protective microbiota function depends on community context: stable and compatible communities allow protection to emerge, whereas dysbiotic communities can lose protective capacity despite containing strains with beneficial potential.

## Discussion

Plants depend on their microbiota for growth, resilience, and disease resistance, making microbial community assembly a key determinant of plant health ^60–62^. Using two barley SynComs, we show that microbiota function is shaped not only by strain-level beneficial potential, but by host genotype and microbial interaction structure. Low intra-microbiota antagonism emerged as a central feature of a stable and protective barley root community, whereas strong internal antagonism was associated with dysbiosis, competitive dominance, and impaired host growth.

Host genotype contributed to root microbiota composition and influenced plant responses to microbial inoculation. Consistent with previous studies, the cultivated barley genotype GP and the wild accession HID4 harbored distinguishable root microbiota despite broad similarities in overall diversity and taxonomic composition ^31,32^. This supports the concept that host genotype contributes to microbiota filtering within barley ^63–66^. However, SynCom inoculation revealed an additional layer of specificity: *Hv*SC1 protected both GP and HID4 against *Bs*, but reduced root growth only in HID4. Thus, microbiota-mediated protection can be maintained across barley genotypes, whereas growth compatibility appears more host-genotype dependent.

A major implication of our work is that strain-level beneficial potential is insufficient to forecast community-level function. *Hv*SC2 was assembled from strains affiliated with genera previously associated with plant growth promotion or pathogen suppression, and many of its members inhibited fungal pathogens *in vitro*. Nevertheless, this community became unstable, failed to protect barley roots, and impaired plant growth. This illustrates a key limitation of trait-prioritized SynCom design: traits beneficial in isolation may be lost, masked, or even become detrimental in combination. In contrast, *Hv*SC1, assembled from frequent members of the natural barley root microbiota, showed lower internal antagonism, greater stability, and host-compatible protection. Natural co-occurrence may therefore capture ecological compatibility among strains and serve as a useful criterion for designing stable microbial communities.

The behavior of *Pseudomonas* strain 397 highlights this context dependence. As an individual strain, 397 showed pathogen-protective capacity, consistent with reports that related *Pseudomonas chlororaphis* isolates can act as biocontrol agents ^59^. In *Hv*SC2, the same strain contributed to competitive dominance and impaired host growth, whereas in *Hv*SC1 it was constrained and did not compromise plant performance. This suggests that microbial traits are not intrinsically beneficial or detrimental, but are shaped by the ecological environment in which a strain operates. For biocontrol applications, highly active strains may be effective only when embedded in communities that buffer their competitive effects and prevent uncontrolled expansion.

Strong negative interactions can destabilize microbial communities by promoting competitive exclusion and reducing diversity, whereas low intra-microbiota antagonism can support coexistence and functional resilience ^15,18,67^. In our system, *Hv*SC1 constrained strain 397 without any single member directly inhibiting it, suggesting that buffering emerged from collective community properties rather than from one dominant antagonist. Potential mechanisms include niche occupation, resource competition, metabolic interdependence, spatial exclusion, or host-mediated filtering. Identifying these mechanisms will be important for understanding how stable microbiota suppress disruptive members while maintaining beneficial functions.

In this experimental comparison, the core SynCom resembling the natural barley root microbiota exhibited greater stability than the trait-prioritized SynCom across multiple complementary readouts. These included sustained strain richness, closer retention of the inoculum composition, preservation of class-level community structure, and higher resistance to dominance by individual competitors (Fig. 6). This higher stability translated into consistent pathogen protection without detrimental effects on host growth in the native genotype. While broader testing across additional SynComs, host genotypes, soils, and assembly strategies will be needed to assess generality, these findings support the idea that natural microbiota, together with the host, favor compatible interaction networks that stabilize both community assembly and function *in planta*.

**Figure 6:**
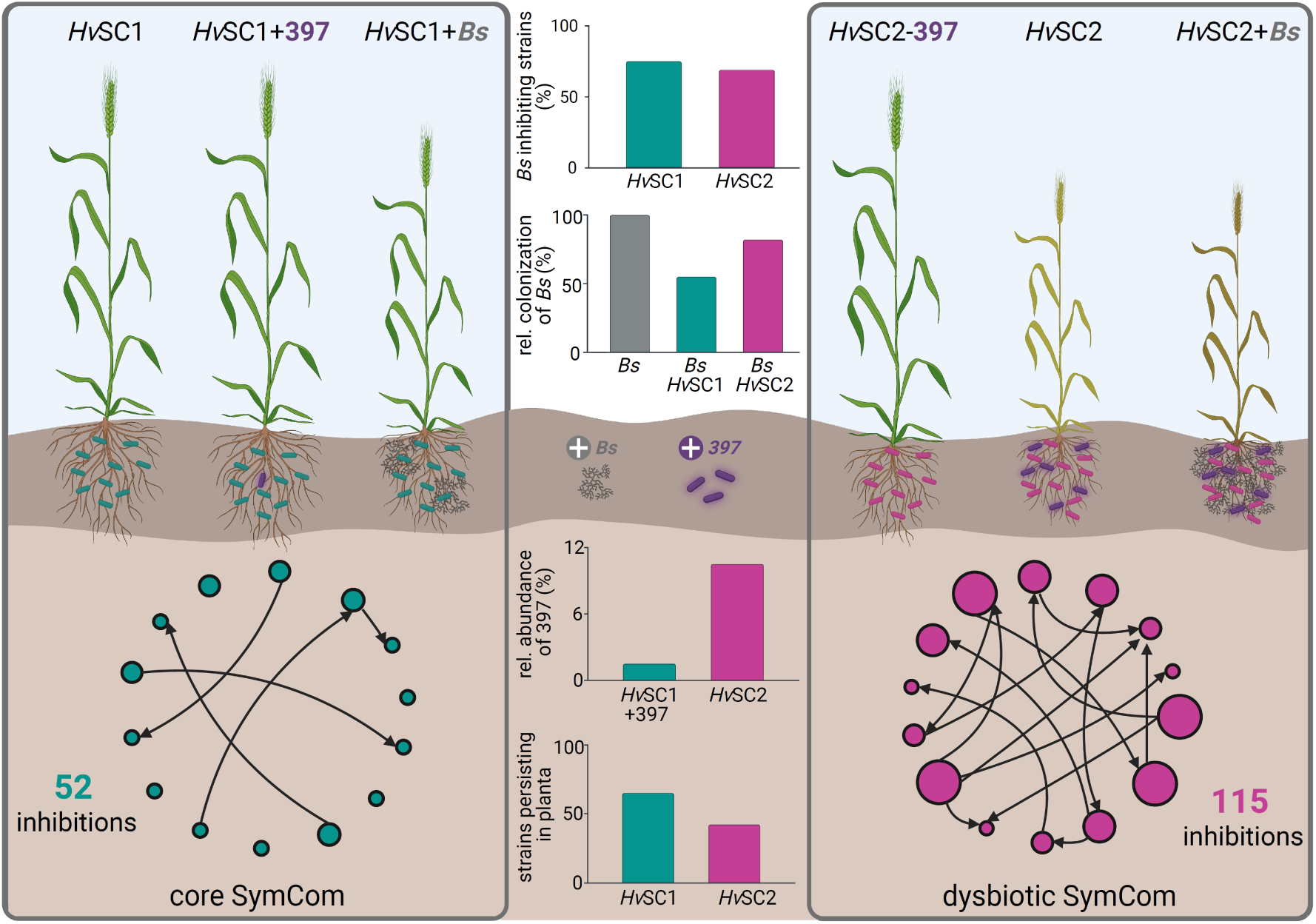
Schematic model of community compatibility, stability, and pathogen protection in barley roots. The core SynCom *Hv*SC1 (left) exhibits fewer antagonistic interactions among bacterial members, with inhibitory interactions indicated by black arrows. This community remains more stable *in planta* and is associated with reduced proliferation of the highly competitive *Pseudomonas* strain 397, as well as reduced colonization by the fungal pathogen *Bipolaris sorokiniana* (*Bs*). *Bs* colonization is further reduced in an inter-kingdom context with the beneficial root endophyte *Serendipita indica* (*Si*). In contrast, the dysbiotic SynCom *Hv*SC2 (right) exhibits more antagonistic interactions, which are associated with reduced community stability. In this context, strain 397 can proliferate and impair barley root growth. Moreover, the protective potential of individual strains within the SynCom is lost at the community level, and *Bs* colonization is not reduced.

Our genomic analyses further show that intra-microbiota antagonism cannot be explained by biosynthetic potential alone. Although BGC richness and phylogenetic relatedness did not explain the greater antagonism in *Hv*SC2, BGC number correlated with halo production in Burkholder plate assays specifically within *Hv*SC2. NRP-metallophore and ranthipeptide clusters were associated with antagonistic activity only in this community. Because these BGC classes were not enriched in *Hv*SC2 relative to *Hv*SC1, their contribution to intra-bacterial community antagonism likely depends not only on their presence, but also on ecological activation, regulation, or metabolite deployment. Thus, biosynthetic potential may translate into antagonism only under particular community conditions. This aligns with recent research on gut bacterial communities, demonstrating that community context is the dominant driver of changes in the microbial proteome ^68^. Future transcriptomic and metabolomic analyses will be required to determine whether specific BGCs are differentially expressed or chemically active in stable versus dysbiotic communities.

The interaction between *Hv*SC1 and the fungal endophyte *Si* extends this concept to inter-kingdom microbiota function. *Hv*SC1 enhanced *Si*-mediated protection against *Bs* without compromising *Si* colonization, indicating compatibility between beneficial bacterial and fungal members in the barley root environment. This suggests that stable bacterial communities can restrain pathogenic fungi while preserving beneficial fungal associations. The basis for this selectivity remains largely unresolved, but may involve pathogen-specific antagonism, immune modulation, spatial niche partitioning, or metabolite-mediated interactions.

Together, our findings suggest that effective plant microbiome engineering should move beyond maximizing individual beneficial traits. Communities resembling natural assemblies may better preserve interaction structures that support coexistence, constrain aggressive competitors, and maintain host protective function. In contrast, trait-prioritized communities may combine strains selected for predicted beneficial traits in ways that generate excessive competition and dysbiosis. Low intra-microbiota antagonism therefore provides a practical and testable principle for designing protective plant-associated microbial communities.

In conclusion, protective barley root microbiota are defined not only by predicted beneficial traits of their members, but also by community compatibility, likely reflecting host filtering and stable, long-standing coexistence. Thus, prioritizing coexistence, stability, and host compatibility alongside beneficial strain traits may improve the design of microbiome-based strategies for sustainable crop protection.

## Material and Methods

### Plant, fungal, and bacterial materials

Barley (*Hordeum vulgare* cv Golden Promise, GP) and the natural barley line *H. vulgare ssp. spontaneum* HID4 was used in this study ^69^. Root amplicon data was further generated from a variety of barley accessions belonging to the species *Hordeum euclaston* (NGB8535, NGB90233, NGB90342), *Hordeum pubiflorum* (NGB6576, NGB6583), *Hordeum multicum* (NGB6545, NGB6547, NGB90357), *Hordeum bogdanii* (*NGB8519, NGB90005, NGB90046)*, *Hordeum vulgare* (Golden Promise, Igri, Morex) and *Hordeum vulgare subsp. spontaneum* (HID4, HID64, HID369). *Bipolaris sorokiniana* (*Bs*, ND90Pr, obtained from S. Zhong, Cereal Disease Laboratory Saint Paul, USA), *Serendipita indica* (*Si*, DSM11827, German Collection of Microorganisms and Cell Cultures GmbH, Braunschweig, Germany) and *Fusarium graminearum* (*Fg*, JLU Giessen, Germany), were used as fungal models. The bacteria used were isolated from the roots of barley GP and HID4 plants grown in natural CAS soil ^70^ for 8 weeks at a day/night cycle of 16/8 h at 22/18 °C, 60% humidity under 108 µmol/m^2^s light intensity. Bacterial isolation was performed as previously described^71^. The bacteria used in this study are listed in Table S7.

### Growth conditions and microbe inoculations

Barley seeds were surface sterilized in 6% sodium hypochlorite for 1 h under continuous shaking and subsequently rinsed repeatedly with sterile water at 30 min for intervals for 4 h. The seeds were germinated on wet filter paper in darkness at room temperature for 4 days, transferred to 1/10 PNM (Plant Nutrition Medium, pH 5.7) ^72^ in sterile glass jars for growth at a day/night cycle of 16/8 h at 22/18 °C, 60% humidity under 108 µmol/m^2^s light intensity. *Si* cultures were propagated on complete medium (CM) ^73^ supplemented with 2 % (w/v) glucose and 1.5 % (w/v) agar at 28 °C in darkness for 28 days prior to spore isolation. *Bs* was propagated on modified CM medium ^74^, containing 1.5% agar at 28 °C in darkness for 14 days prior to inoculation. *Fg* was propagated on PDA medium containing 1.5% agar at 28 °C in darkness for 14 days pre inoculation. *Si* spores and *Bs* conidia suspensions were prepared as described in^74^. *Fg* spore suspension was prepared as for Bs. Multipartite inoculation assays were performed as described in ^75^.

Single bacterial strains were streaked out and grown on solid TSB medium (Sigma Aldrich) (15 g/l) at 28 °C. Single colonies were picked and grown in liquid TSB medium with shaking at 120 rpm for 1−3 days depending on growth rates. The cultures were 3 times washed with sterile water, each followed by a centrifugation at 3500 rpm for 10 min at 4 °C, to remove the TSB medium. Final OD_600_ was adjusted to 0.01 prior to inoculation of single strains or mixtures in equal amounts for SynCom compositions to a final OD_600_ of 0.01. Barley roots were inoculated with *Bs* conidia (5 × 10^3^ spores/ml), single bacterial isolates (OD_600_ = 0.01), *Si* mycelium (0.04 g/ml), *Fg* spores (5000 spores/ml) or a respective mixture of organisms per jar. Sterile MilliQ water was used as a control treatment. The drop-in- and drop-out experiment were performed using a variety of concentrations of the Pseudomonas strain 397 which all resulted in comparable barley growth phenotype. The data presented in this paper reflect experiments where the Pseudomonas strain 397 was either added to *Hv*SC1 at a final OD_600_ of 0.01 or left out of *Hv*SC2. Barley roots were harvested at 6 days post inoculation (dpi). Per biological replicate of each experiment and treatment, roots from four barley plants were pooled. For RNA extraction barley roots were washed thoroughly to remove extraradical fungal hyphae and epiphytic bacteria and snap-frozen in liquid nitrogen.

### RNA isolation for RNA-seq and RT-PCR

RNA extraction for quantification of fungal endophytic colonization, cDNA generation, and RT-PCR were performed as described previously ^74^. The primers used are listed in Table S8.

### Modified Burkholder agar diffusion assay

For studying inhibitory interactions within the bacterial isolates, we modified the Burkholder agar diffusion assay^76^. The bacterial strains were grown and washed as described above. 50 ml of molten agar (42 °C) was mixed with 600 μl of the target bacterial strain (OD_600_ = 1) and poured onto a square plate (12cm x 12 cm). After the agar solidified, 10 μl drops of nine potential producer strains (OD_600_ = 0.1) were placed onto the plate in a 3 x 3 matrix. The plates were incubated at 28 °C in darkness for 4 days. Afterwards the presence of and inhibition zone (an area in which the target isolate failed to growth, hereafter referred to as a halo) was assessed. Inhibition networks were visualized in R using the package igraph v2.2.1 ^77^.

### Fungal inhibition assay

To test the bacterial strains for their ability to inhibit fungal growth, we performed confrontation assays on plate. 1 ml *Bs* conidia (5 x 10^3^ spores/ml) or 1 ml *Si* spores (5 x 10^6^ spores/ml) were spread on solid TSB medium, either alone or mixed with individual bacterial isolates (OD_600_= 0.1). The plates were incubated at 28 °C for up to 6 days. Fungal growth was determined using a stereomicrosope. Fungal inhibition was categorized in three groups: no inhibition, partial inhibition, full inhibition. Full inhibition was determined by an inhibition of or shortly after spore germination. Partial inhibition was categorized by a lower growth compared to the fungus grown alone. For high resolution microscopy, 1 ml *Bs* conidia (5 x 10^3^ spores/ml) were spread onto solid 1/10 PNM medium, either alone or in combination with *Pseudomonas* strain 397 (OD_600_ = 0.01). Thin sections of the agar surface were excised using a sterile scalpel and subjected to imaging. Microscopy was performed using a Leica THUNDER widefield fluorescence microscope (Leica Microsystems GmbH, Wetzlar, Germany).

### Barley root natural community profiling und culture collection characterization

Barley root bacterial communities were profiled using amplicon sequencing of the V5-V7 regions of the bacterial 16S rRNA gene (Table S8). For GP and HID4, plants were grown in CAS soil as described above, the additional Barley species and accessions were grown in a CAS soil/sand mixture in a common garden experiment and harvested 45 days post sowing. All roots were harvested, washed in ice-cold water and surface sterilized as described above. DNA was extracted using the NucleoSpin DNA extraction Kit (Macherey-Nagel). All amplicon sequencing was done using the Illumina MiSeq platform, either at the Max Planck Genome Centre, Cologne, Germany, or at the Cologne Centre for Genomics CCG, Germany.

In order to cross-reference the natural community and culture collection OTUs, BLAST v2.5.0+ ^78^ was used. The natural community OTUs (NatCom_OTU) remaining after filtering were compared to the culture collection OTUs (CultColl_OTU), ignoring end-gaps, to account for differential trimming depending on the sequencing run quality. The BLAST results were filtered to retain only the highest-identity match for each NatCom_OTU. A NatCom_OTU was considered ‘recovered’ in the culture collection if it showed a ≥97% identity match to a CultColl_OTU derived from the same barley accession. The recovery rate was then calculated for each barley accession, as the percentage of recovered OTUs from the top 100 most abundant NatCom_OTUs (*i.e.*, with the highest mean relative abundance). The accession-specific cumulative recovered relative abundance was defined as the sum of the mean relative abundance of each recovered NatCom_OTU (Fig. S1C-F).

### Amplicon sequencing data analysis

For comparability, amplicon sequencing data processing from both the NatCom profiling and the CultColl isolate identification was largely done with the same pipeline. NatCom samples yielded at least 150000 paired-end 300 bp reads per sample, whereas CultColl isolates yielded approximately 5000 paired-end 300 bp reads per sample. Demultiplexed amplicon sequencing data were pre-processed with Trim Galore v0.6.10 to filter and trim low-quality reads, remove primer and adapter sequences ^79^. The quality of the raw and filtered reads was assessed using FastQC v0.12.1 ^80^ and multiQC v1.3.0 ^81^. Follow-up processing steps were performed using the vsearch v2.24.0 framework ^82^. Forward and reverse reads were merged and filtered, and the resulting sequences were de-replicated, denoised (UNOISE3), and chimeras were detected *denovo* and removed with the UCHIME3 algorithm. Merged reads were quality-filtered by removing sequences containing ambiguous bases, sequences shorter than 225 bp, and reads exceeding a maximum expected error threshold of 0.5. Full-length de-replication, denoising and chimera removal were performed on a per-sample basis prior to global sequence pooling. For one sequencing run of slow-growing isolates that yielded higher overall sequence quality and at least 5000 paired-end 300 bp reads per sample, denoising stringency was increased by lowering the UNOISE alpha parameter from the default value of 2.0 to 0.8.Additionally, representative sequences from this run were clustered at 99% nucleotide identity prior to concatenation with the remaining culture collection sequences, to account for the elevated sequence resolution observed in that dataset.

Following per-sample processing, all non-chimeric sequences were pooled and clustered at 100% nucleotide identity to generate zero-radius operational taxonomic units (zOTUs). The 99% identity clusters from the slow-growing isolate run described above were then used for cross-run integration. Representative sequences from all culture collection sequencing runs were subsequently concatenated and re-clustered at 100% identity to ensure consistent operational taxonomic unit definitions across datasets. The resulting zOTUs were taxonomically classified against the RDP v19 database ^83^ using the non-Bayesian taxonomic classifier SINTAX (--sintax_cutoff 0.6), within the vsearch v2.28.1 framework. Prior to taxonomic assignment, low-abundance and likely erroneous sequences were reduced through UNOISE denoising and de novo chimera removal.

Amplicon sequencing data from *Hv* roots inoculated with *Hv*SC1 and *Hv*SC2 including drop-in and drop-out of strain 397 after 6 dpi were processed slightly differently due to lower complexity compared with natural soil samples. Raw Illumina paired-end 250 bp reads (approximately 50000 read pairs per sample) were first subjected to adapter and primer trimming using cutadapt v5.1 ^84^ followed by quality-based trimming (Phred score threshold 27) and the removal of reads containing ambiguous bases or shorter than 100 bp. Quality control at each preprocessing step was performed using FastQC v0.12.1 and MultiQC v1.12. Read pairs were merged using flash2 v2.2.00 ^85^ with default parameters suitable for overlapping amplicons.

### Natural community and culture collection characterization

The natural community and culture collection data was subsequently analyzed in R v 4.4.2, where the different steps (relative abundance calculation, alpha-diversity, beta-diversity, unconstrained and constrained dimensionality reduction) were performed with the R package vegan ^86^. Differential abundance analysis was performed using DESeq2, and adjusted effect sizes were calculated with the apeglm method ^87^. The identification of the core microbiome was done as previously described, using a binomial distribution to determine presence and/or absence probability ^88^. Phylogenetic trees of the natural community members were constructed using the maximum likelihood method, with IQ-TREE v2.0.3 ^89^ and QIIME2 v2020.8.0 ^90^ based on a masked alignment (MAFFT v 7.471 ^91^ Biostrings v2.74.1 ^92^was used to manage FASTA sequences in R. Phylogenetic tree handling and vizualization was performed in R, using ape v5.8-1 ^93^, TreeTools v2.0.0 ^94^, ggtree v3.14.0 ^95^ ggtreeExtra v1.16.0 ^96^ and ggnewscale v0.5.2 ^97^.

### Whole genome sequencing, assembly and annotation of SynCom members

The DNA from the selected SynCom members was isolated using the Monarch HMW DNA Extraction Kit (NEB). Quality control, library preparation and sequencing were carried out in collaboration with CCG, using Oxford Nanopore Technologies on MinION and PromethION flow cells.

Raw Nanopore reads were basecalled and demultiplexed with the standard CCG pipeline, and assemblies were generated with Flye v2.8.3 ^98^ using the --nano-raw mode. Due to high estimated sequencing coverage (>1500×), all assemblies were performed with internal coverage downsampling (-asm-coverage 60) and a conservatively estimated genome size of 9 Mb to prevent redundancy-induced graph collapse. Assembly quality was assessed using multiple complementary tools. QUAST v5.3.0 ^99^ was applied to evaluate contiguity, total length, and N50 values. To detect potential mis-assemblies, structural inconsistencies, and uncorrected errors, Inspector v1.3 ^100^ was used. Assemblies were polished using Inspector’s correction module, and corrected contigs were re-evaluated in a second Inspector run. Only corrected contigs passing Inspector’s validity criteria were retained for downstream analyses. To assess genome completeness and detect possible contamination, each final assembly was analyzed with BUSCO v6.0.0 ^101^ using the bacteria_odb10 lineage dataset (Table S9). Taxonomic screening was performed with Kraken2 v2.1.3 ^102^, using the default database and contaminated contigs were removed prior to re-evaluation. Only assemblies exhibiting high completeness and low duplication were retained for further analyses. Genome annotation was performed using Bakta v1.11.4 ^103^.

### Functional Genome Analysis

Biosynthetic gene clusters (BGCs) were predicted from genome assemblies using antiSMASH v7.1.0 ^104^ with the bacterial taxon setting and full annotation enabled. All BGC loci identified per strain were extracted and categorized by cluster type. The antiSMASH output (Table S10) was used to construct a per-strain BGC count matrix (strains × BGC types), which was used for all downstream analyses. To group predicted BGCs into gene cluster families (GCFs) based on shared biosynthetic logic, BiG-SCAPE v 2.0.2 ^52^ was run on the full antiSMASH output using the Pfam-A HMM database. GCF clustering was performed in mixed-mode (--mix), including singleton clusters (--include-singletons), with a cutoff threshold of 0.5. The resulting presence/absence matrix (Table S11) of GCFs per strain was used to compute pairwise GCF Jaccard distances between all strains using the vegdist function (method = “jaccard”, binary = TRUE) from the vegan R package.

Genome-based phylogenetic distances were computed using GTDB-Tk (v2.6.1) with the GTDB release226 reference database. The “classify_wf” workflow was applied to all genome assemblies (--skip_ani_screen), producing a multiple sequence alignment (MSA) across 120 conserved bacterial marker genes (bac120). Pairwise p-distances were calculated from this MSA as the fraction of alignment columns differing between each genome pair, excluding gap-only positions.

### Amplicon-based SynCom abundance estimation

To generate a 16S reference database for all SynCom isolates, 16S rRNA gene sequences were extracted from genome assemblies when available. For isolates of which no usable genome assembly could be obtained, the corresponding culture collection OTU sequence was used as a substitute reference (Table S7). First, using bakta generated GFF3 annotation files, all annotated 16S rRNA loci were extracted in full length from each genome. To ensure compatibility with the 16S amplicon region used in the experimental sequencing (V5–V7 region), the full-length sequences were next trimmed in silico using the same primer pair as in the wet-lab protocol (Table S8). Using the full 16S rRNA sequences as reference, primer matching was performed with cutadapt v5.1, allowing up to two mismatches per primer and enforcing a three-base exact match at the 3′ end to reflect the moderate stringency of the PCR annealing temperature used in the experiment. Only amplicons between 100 and 500 bp were retained. For each isolate, amplicons were deduplicated so that only unique 16S sequence variants were kept. All unique amplicons across isolates were finally compiled into the SynCom-specific 16S reference databases used for downstream mapping. To normalize the abundance data for reference copy number, multiple appearances of identical sequences were recorded as copy number.

Preprocessed, merged reads of amplicon sequencing data were processed using a custom pipeline based on vsearch v2.30.1. All samples were first dereplicated using vsearch --fastx_uniques. Dereplicated reads were then mapped against the genome-derived SynCom 16S reference databases using vsearch --usearch_global, requiring a minimum sequence identity of 97% and a minimum query coverage of 95%. All candidate alignments were retrieved per read (--maxaccepts 0, --maxhits 0) to enable downstream disambiguation. Mapping was performed in two hierarchical passes: in the first pass, reads were mapped against the reference sequences of isolates expected in the respective sample type (priority references). Reads that did not match any priority reference were then mapped in a second pass against all remaining references. Ambiguous mappings were defined as reads with multiple equally ranked hits at both identity and query coverage and were flagged and excluded from downstream quantification. Raw count tables were generated from unambiguous mappings only.

To remove spurious detections, a biological filtering step was applied per sample type. Within each treatment group, a reference sequence was retained only if it was detected in at least two replicates and had a minimum of 10 total mapped reads. Copy-number correction was performed by dividing raw read counts by the 16S rRNA gene copy number assigned to each reference sequence. After copy-number normalization, abundances belonging to different 16S variances of the same isolate were aggregated into a single isolate-level abundance estimate.

All downstream analyses and visualization, including aggregation of abundance tables, calculation of means, relative abundances, and alluvial plots were performed in R v4.4.2 utilizing the tidyverse v2.0.0 ^105^ and ggalluvial v0.12.5 ^106^ packages.

#### Statistical analyses

For colonization, plant phenotypes, disease symptoms, and gene expression, data were tested for normality and homogeneity of variances. Depending on the experiment and number of factors, statistical analyses were performed using one-way or two-way ANOVA followed by Tukey’s post-hoc test (p < 0.05), or the non-parametric Kruskal–Wallis test when assumptions were not met. If the Kruskal–Wallis test indicated significant differences, pairwise comparisons were performed using Dunn’s test with FDR-adjusted p-values (< 0.05). For direct comparisons between two groups, a two-sample Student’s t-test was applied when assumptions of normality and homoscedasticity were fulfilled; otherwise, the non-parametric Mann–Whitney U test was used. For BGC analyses, associations between categorical groupings (e.g. inhibition level or halo production status) and BGC type prevalence were assessed using contingency-based tests on per-strain presence/absence data, with multiple testing correction applied using the Benjamini-Hochberg false discovery rate procedure. The relationship between total BGC repertoire size and inhibitory activity was assessed by correlation, and the contribution of individual BGC types to halo production was evaluated using linear models and non-parametric rank-based tests. To investigate whether phylogenetic relatedness and genomic biosynthetic similarity predict inhibitory interactions between strains, distance-based correlation analyses were performed on pairwise phylogenetic, GCF content, and inhibition profile distance matrices. All p-values from multiple comparisons were corrected for false discovery rate. Specific tests and parameters are indicated in figure legends.

## Acknowledgements

We thank Professor Maria von Korff for providing plant material from additional barley accessions for root amplicon sequencing, and Dr. Ruben Garrido-Oter for access to the Illumina MiSeq platform at the MPIPZ. We further thank Tania Matsumoto for her support to the sample preparation for amplicon sequencing. LM, NP, SA, GA, GL, and AZ acknowledge support from the FOR 5682 MadFungi, funded by the Deutsche Forschungsgemeinschaft (DFG, German Research Foundation) under Project ID 520490591 and the SPP2125 DECRyPT. SK and AZ acknowledge support from the Cluster of Excellence on Plant Sciences (CEPLAS), funded by the DFG under Germany’s Excellence Strategy - EXC 2048/2 - Project ID 390686111.

## Author contributions

LM, GL, and AZ conceived and designed the study and conceptualized the manuscript. LM and GA prepared samples for characterization of the natural barley root microbiota and performed laboratory work related to fungal and bacterial inoculation assays. LM, GA, and SPK performed fungal and bacterial confrontation assays. SPK performed microscopy. SA and LM designed the synthetic communities, and SA established the culture collection using bioinformatic approaches. GA maintained and organized the culture collection. NP analyzed amplicon-based SynCom abundance data and performed functional genomic characterization. SA and NP conducted genome assembly. AZ secured funding for the project. All authors contributed to manuscript editing and approved the final version.

## Declaration of generative AI and AI-assisted technologies

The authors used ChatGPT (OpenAI, San Francisco, CA, USA) to support language refinement, editing, and clarity improvements during manuscript preparation. Perplexity AI (Perplexity AI, Inc., San Francisco, CA, USA) was used to support literature checking and source verification. AI-assisted tools were not used to generate original scientific hypotheses, data, analyses, or conclusions. All AI-assisted content was critically reviewed, edited, and approved by the authors, who take full responsibility for the final content of the manuscript.

## Data availability

All data supporting the findings of this study are available within the article and its Supporting Information and will be made available through the ARC repository under DOI [ARC DOI]. Genome assemblies of the bacterial isolates generated in this study will be deposited in NCBI GenBank under accession number [accession number].

