## Supplemental Figures for "Low intra-microbiota antagonism underlies stable and protective barley root microbiota"

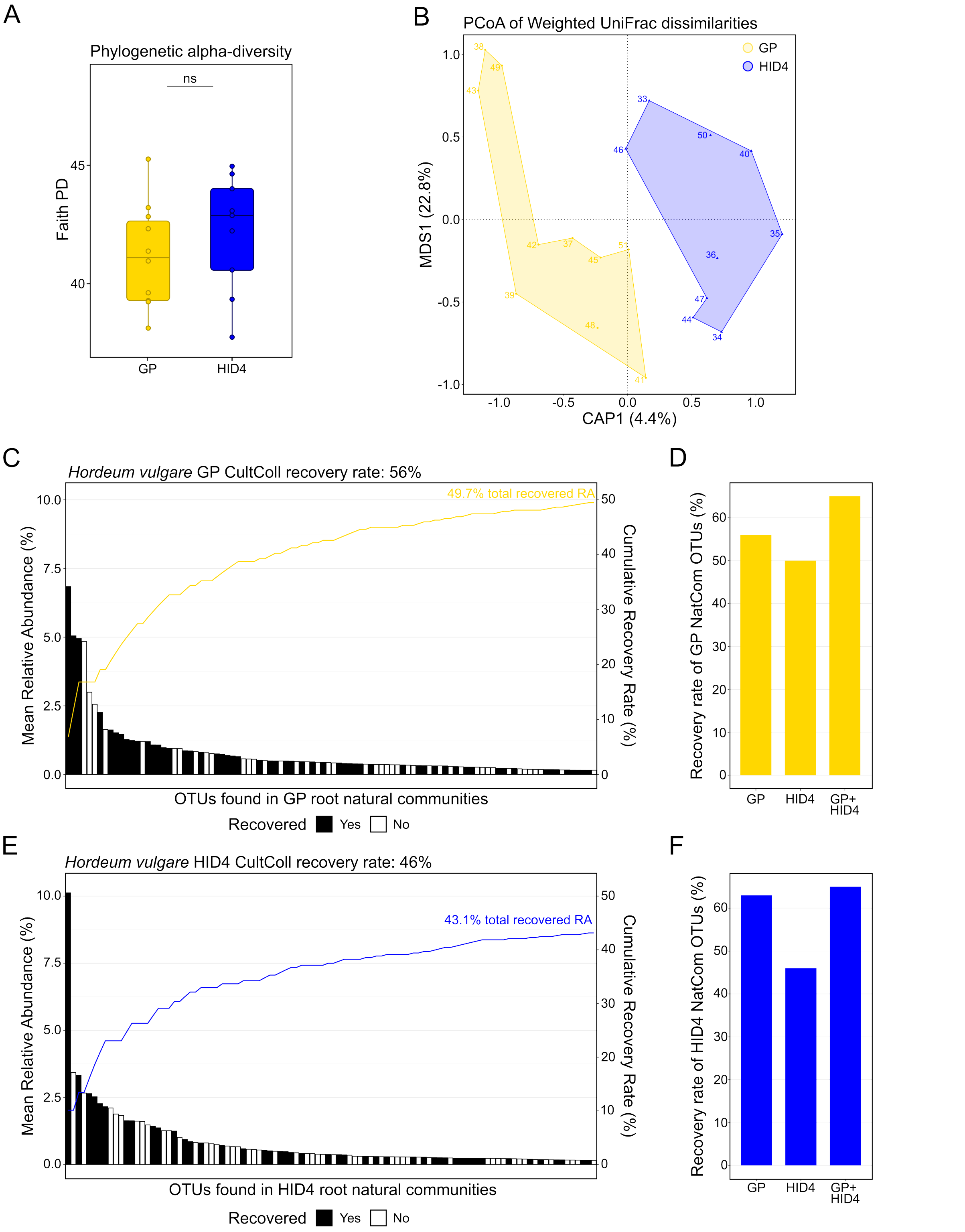


**Figure S1: Natural community alpha- and beta-diversity and recovery rate.** A) Faith's Phylogenetic Diversity (Faith's PD) of root bacterial communities from GP (gold) and HID4 (blue) barley. Box plots show the distribution across biological replicates per genotype. Statistical significance between genotypes was assessed with a Wilcoxon rank-sum test. B) Principal Coordinates Analysis (PCoA) of weighted UniFrac dissimilarities between root bacterial communities of GP and HID4 barley, accounting for both phylogenetic relatedness and relative abundance. Each point represents one biological replicate; shaded polygons indicate group membership (GP: gold; HID4: blue). The percentage of total variance explained by each axis is indicated in parentheses. Differences in community composition between genotypes were tested by PERMANOVA (9,999 permutations), with homogeneity of group dispersions confirmed by permutation test on multivariate dispersions (betadisper/permutest, 9,999 permutations). C) and E) Recovery of natural community OTUs in the culture collection for GP and HID4, respectively. The 100 most abundant natural community (NatCom) OTUs are ordered on the x-axis by decreasing mean relative abundance. Bars show the mean relative abundance of each NatCom OTU, and bar fill indicates whether the OTU was recovered in the culture collection at ≥97% nucleotide identity. The line (gold for GP and blue for HID4) shows the cumulative recovered relative abundance. D) and F) Total number of top 100 NatCom OTUs recovered at ≥97% identity for GP and HID4, respectively. On the x-axis, recovered counts are grouped by the genotype from which culture-collection isolates were obtained: GP, HID4, or the combined GP and HID4 collection. Culture collection sequences were generated from isolates obtained independently from GP and HID4 root samples; pairwise sequence comparisons were performed using BLASTN, retaining the top hit per query.

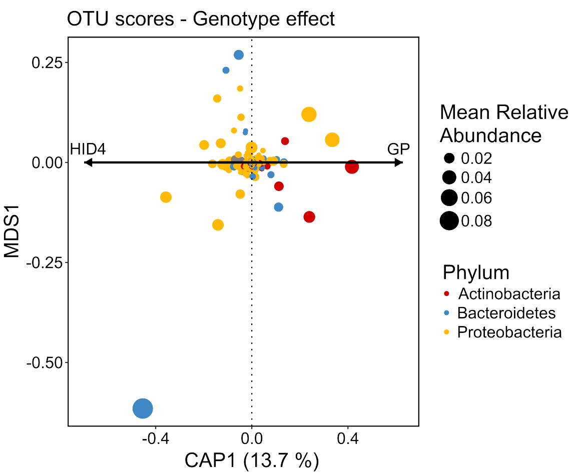


**Figure S2: Taxonomic drivers of genotype-dependent root microbiota differentiation in GP and HID4 barley roots.** Distance-based redundancy analysis (dbRDA) ordination of Bray–Curtis dissimilarities was used to assess the effect of genotype on bacterial community composition. Arrows indicate the direction of genotype-associated vectors for GP and HID4 in ordination space. Overlaid OTU scores represent taxa associated with genotype-dependent separation, colored by phylum and scaled by mean relative abundance.

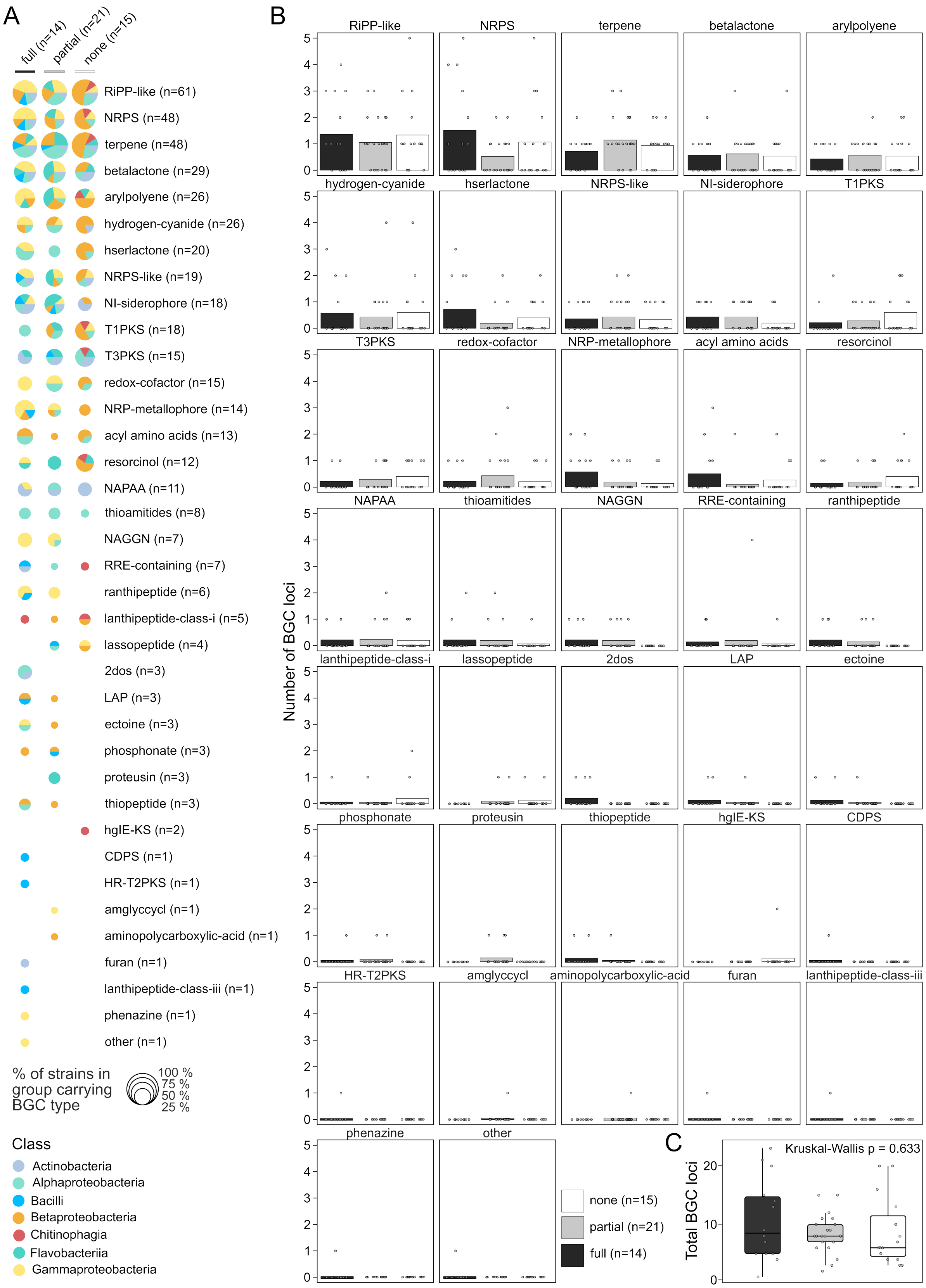

A
full (n=14)
partial (n=21)
none (n=15)
RiPP-like (n=61)
NRPS (n=48)
terpene (n=48)
betalactone (n=29)
arylpolyene (n=26)
hydrogen-cyanide (n=26)
hserlactone (n=20)
NRPS-like (n=19)
NI-siderophore (n=18)
T1PKS (n=18)
T3PKS (n=15)
redox-cofactor (n=15)
NRP-metallophore (n=14)
acyl amino acids (n=13)
resorcinol (n=12)
NAPAA (n=11)
thioamitides (n=8)
NAGGN (n=7)
RRE-containing (n=7)
ranthipeptide (n=6)
lanthipeptide-class-i (n=5)
lassopeptide (n=4)
2dos (n=3)
LAP (n=3)
ectoine (n=3)
phosphonate (n=3)
proteusin (n=3)
thiopeptide (n=3)
hglE-KS (n=2)
CDPS (n=1)
HR-T2PKS (n=1)
amglyccycl (n=1)
aminopolycarboxylic-acid (n=1)
furan (n=1)
lanthipeptide-class-iii (n=1)
phenazine (n=1)
other (n=1)
% of strains in group carrying BGC type
100 %
75 %
50 %
25 %
Class
Actinobacteria
Alphaproteobacteria
Bacilli
Betaproteobacteria
Chitinophagia
Flavobacteriia
Gammaproteobacteria
B
Number of BGC loci
RiPP-like
NRPS
terpene
betalactone
arylpolyene
hydrogen-cyanide
hserlactone
NRPS-like
NI-siderophore
T1PKS
T3PKS
redox-cofactor
NRP-metallophore
acyl amino acids
resorcinol
NAPAA
thioamitides
NAGGN
RRE-containing
ranthipeptide
lanthipeptide-class-i
lassopeptide
2dos
LAP
ectoine
phosphonate
proteusin
thiopeptide
hglE-KS
CDPS
HR-T2PKS
amglyccycl
aminopolycarboxylic-acid
furan
lanthipeptide-class-iii
phenazine
other
none (n=15)
partial (n=21)
full (n=14)
C
Kruskal-Wallis p = 0.633
Total BGC loci

**Figure S3: BGC repertoire does not explain *Bs* inhibition capability.** A) Scatter pie chart depicting the secondary metabolite biosynthetic potential of SynCom bacteria stratified by their capacity to inhibit *Bipolaris sorokiniana* (*Bs*) growth: full, partial, or no inhibition. BGCs were predicted using antiSMASH v7.1.0, and BGC types are ranked by their total occurrence across all strains. Pie size represents the percentage of strains within each inhibition group carrying a given BGC type; pie color composition reflects the taxonomic class breakdown of carrier strains within each group. For each BGC type, a Fisher's exact test was performed on the 2 × 3 contingency table of inhibition category against carrier/non-carrier status, with p-values corrected for multiple testing using the Benjamini–Hochberg (BH) method. Monte Carlo simulation (B = 10,000) was used due to small expected cell counts in some groups. B) Number of BGC loci for each BGC type per strain, stratified by Bs inhibition capability. Each facet represents one BGC type; bars show group means and grey dots represent individual strains. For each BGC type independently, a Kruskal–Wallis test was performed on per-strain locus counts, including strains with zero counts, followed by pairwise Dunn's post-hoc tests with BH-adjusted p-values. C) Total number of BGC loci per strain across inhibition groups. Grey dots represent individual strains, with the total number of BGC loci detected in each genome shown on the y-axis. Group differences were assessed using a Kruskal–Wallis test.

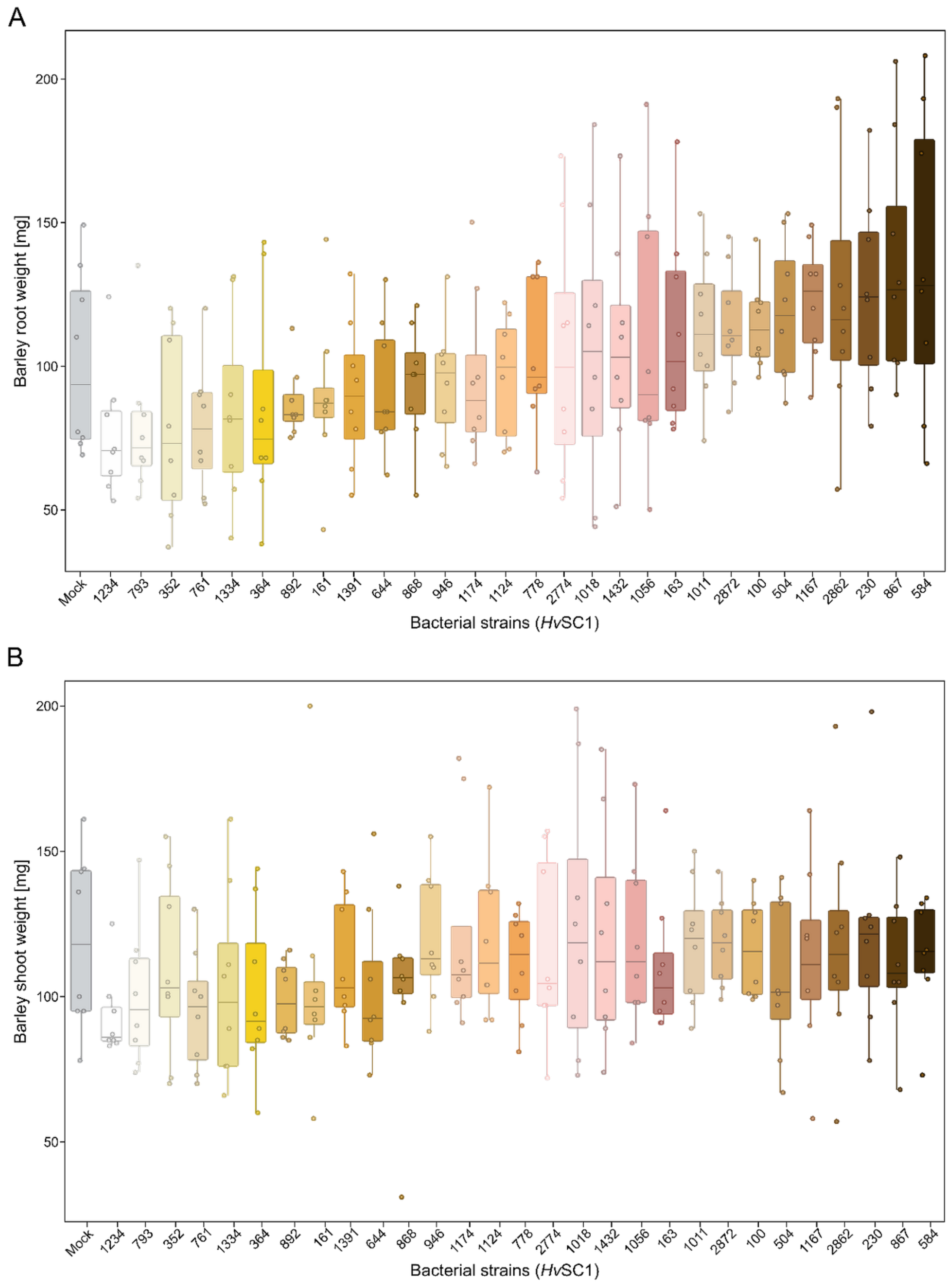


**Figure S4: Barley growth responses to individual *Hv*SC1 strains.** Root and shoot fresh weight of barley cultivar GP in mono-association with individual bacterial strains from *Hv*SC1 at 6 dpi. Box plots show the distribution across biological replicates, with individual points representing replicate values. Statistical differences were assessed using two-sample t-tests against mock ($p < 0.05$).

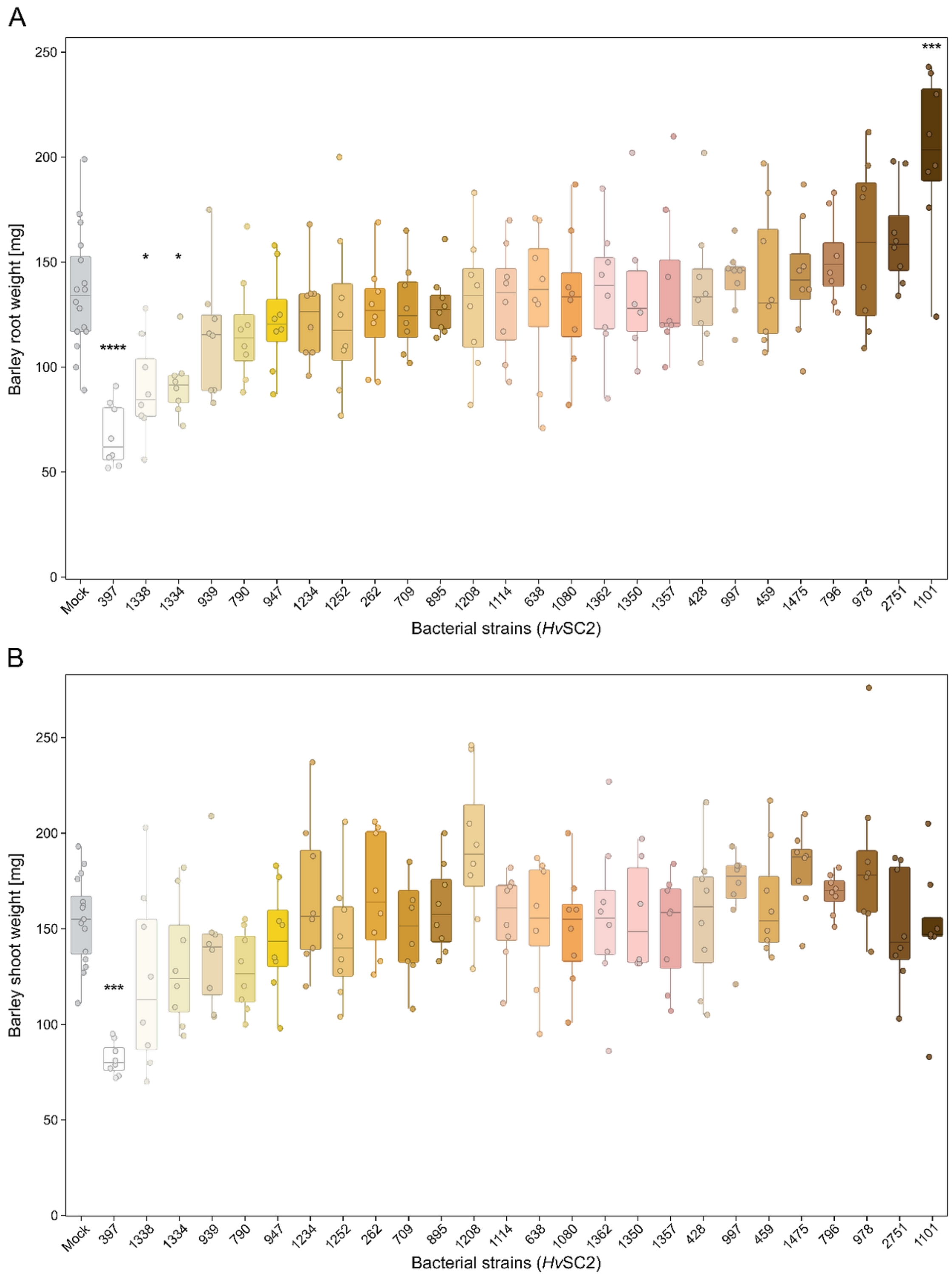


**Figure S5: Barley growth responses to individual *Hv*SC2 strains.** Root and shoot fresh weight of barley cultivar GP in mono-association with individual bacterial strains from *Hv*SC2 at 6 dpi. Box plots show the distribution across biological replicates, with individual points representing replicate values. Statistical differences were assessed using two-sample t-tests against mock ($p < 0.05$).

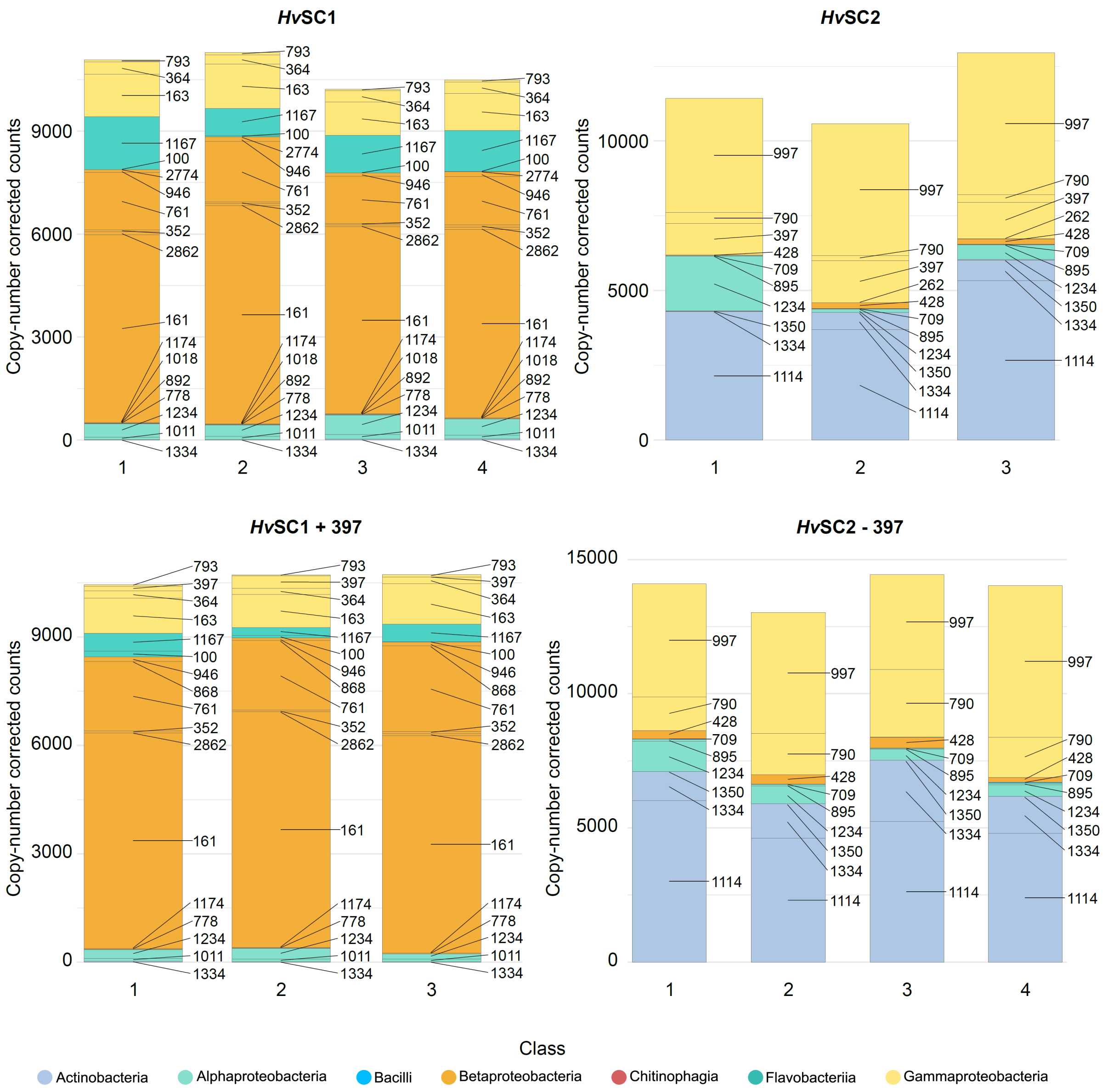


**Figure S6: Total abundance of SynCom isolates per biological replicate across all four experimental treatments at 6 dpi.** Stacked bar plots show the copy-number-corrected absolute abundance of each bacterial isolate for individual biological replicates of *Hv*SC1, *Hv*SC2, Drop-in, and Drop-out treatments after 6 dpi on barley roots. Each bar segment represents one isolate, stacked and coloured by bacterial class. The y-axis shows total mapped read abundance after copy-number correction. Only isolates belonging to the respective SynCom composition are shown.

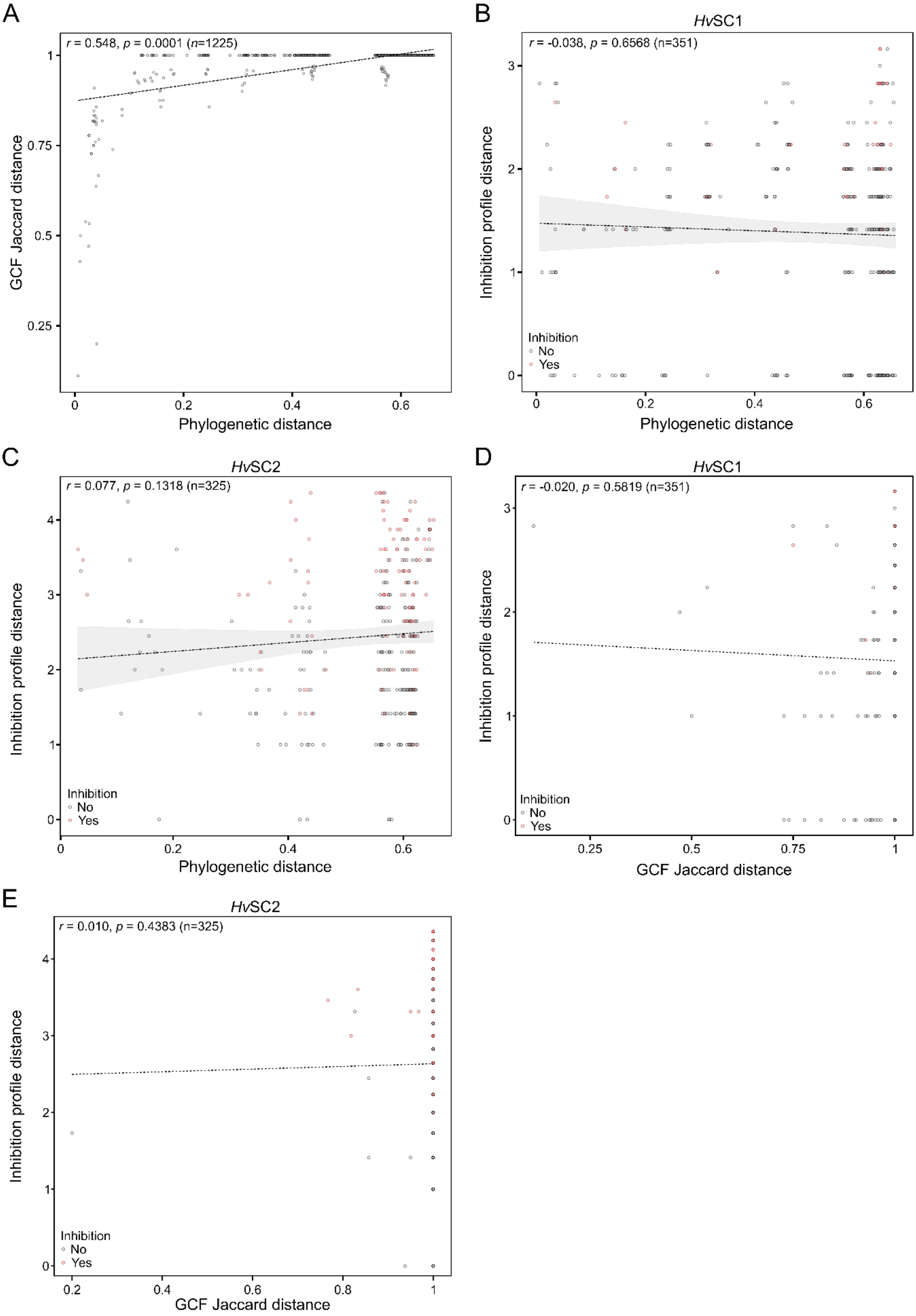
A
r = 0.548, p = 0.0001 (n=1225)
GCF Jaccard distance
Phylogenetic distance
B
HvSC1
r = -0.038, p = 0.6568 (n=351)
Inhibition profile distance
Phylogenetic distance
Inhibition
No
Yes
C
HvSC2
r = 0.077, p = 0.1318 (n=325)
Inhibition profile distance
Phylogenetic distance
Inhibition
No
Yes
D
HvSC1
r = -0.020, p = 0.5819 (n=351)
Inhibition profile distance
GCF Jaccard distance
Inhibition
No
Yes
E
HvSC2
r = 0.010, p = 0.4383 (n=325)
Inhibition profile distance
GCF Jaccard distance
Inhibition
No
Yes

**Figure S7: Phylogenetic distance predicts BGC content similarity, but neither phylogenetic distance nor BGC similarity predicts inhibitory interactions. A)** Pairwise phylogenetic distance, based on bac120 p-distance, plotted against pairwise GCF content dissimilarity, based on Jaccard distance of binary BiG-SCAPE GCF presence/absence, for all strain pairs. A linear regression line is overlaid. The plot is annotated with a permutation-based Pearson correlation coefficient (*r*) and *p*-value based on 9,999 permutations. **B-C)** Pairwise phylogenetic distance plotted against pairwise Euclidean distance of inhibition profiles for *Hv*SC1 (B) and *Hv*SC2 (C). Dot color indicates whether inhibition occurs between the two compared strains: grey = no inhibitory interaction in either direction; red = inhibition in at least one direction. A linear trend line is shown; Mantel *r* and permutation-based *p*-values are annotated. **D-E)** Pairwise GCF distance plotted against pairwise Euclidean distance of inhibition profiles for *Hv*SC1 (D) and *Hv*SC2 (E). Dot color indicates whether inhibition occurs between the two compared strains, as in B and C. A linear trend line is shown; Mantel *r* and permutation-based *p*-values are annotated.

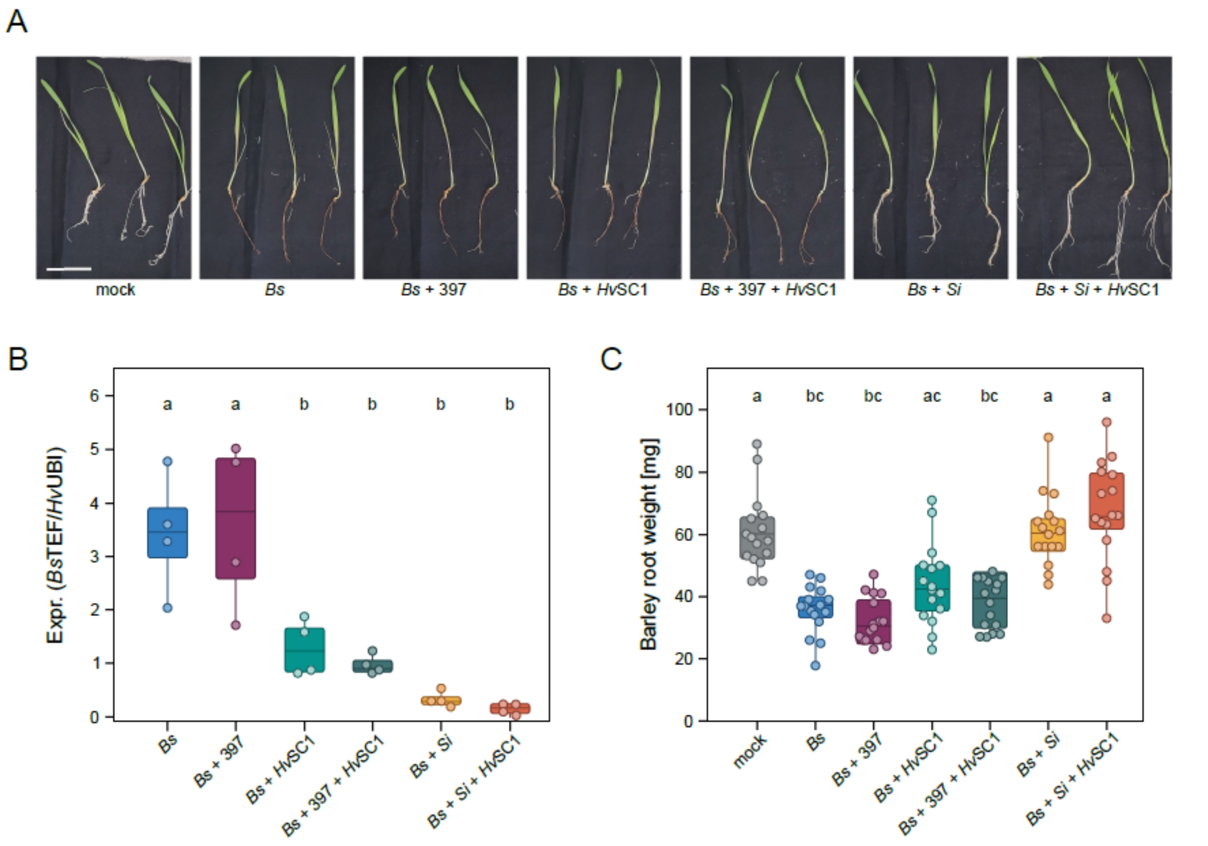


**Figure S8: Bacterial, fungal, and inter-kingdom-mediated host protection in barley accession HID4.** A) Representative images of HID4 roots infected with the fungal pathogen *Bipolaris sorokiniana* (*Bs*) alone or in the presence of *Hv*SC1, *Pseudomonas* strain 397, and the beneficial fungal endophyte *Serendipita indica* (*Si*), either individually or in combination. The white scale bar represents 5 cm. B) Relative colonization of HID4 roots by Bs under the conditions shown in A. C) Barley root fresh weight under the conditions shown in A. Statistical significance was assessed using a Kruskal–Wallis test followed by Dunn's multiple-comparison test ($p < 0.05$).

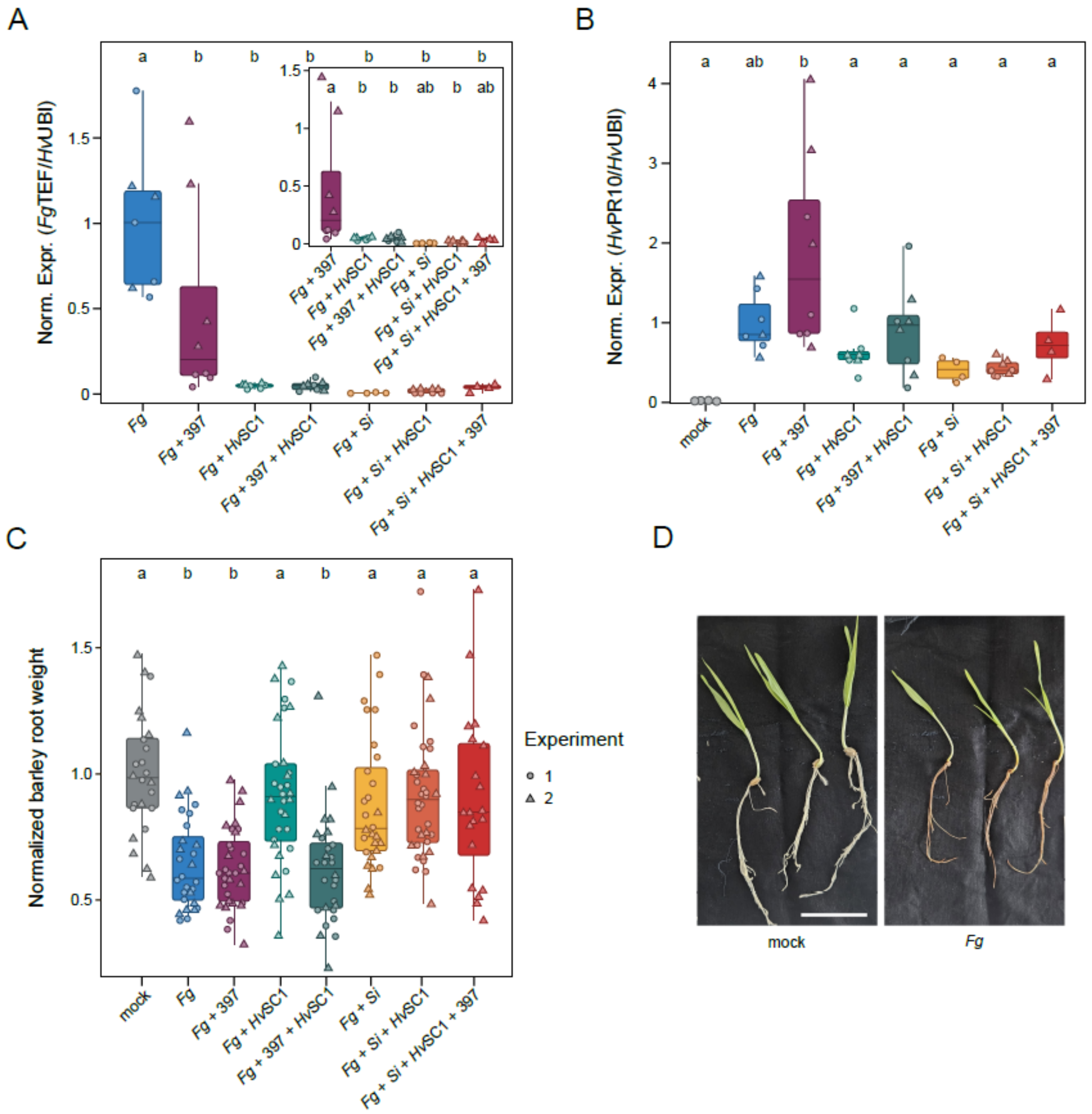


**Figure S9: Bacterial, fungal, and inter-kingdom-mediated host protection against *Fusarium graminearum* in barley cultivar GP.** A) Relative colonization of GP roots by *Fg* inoculated with *Fusarium graminearum* (*Fg*), *Pseudomonas* strain 397, *Hv*SC1, and the beneficial fungal endophyte *Serendipita indica* (*Si*), either individually or in the indicated combinations. Values were normalized to the mean *Fg* colonization level within each experimental round. B) Relative expression of the barley defense marker *PR10* under the conditions described in A. Values were normalized to the mean *PR10* expression under *Fg* treatment within each experimental round. C) Root fresh weight of GP plants under the conditions described in A. D) Representative images of GP roots inoculated with mock or *Fg*. Scale bar, 5 cm. Point shapes indicate independent experimental rounds. Statistical differences were assessed using one-way ANOVA followed by Tukey’s HSD test ($p < 0.05$).

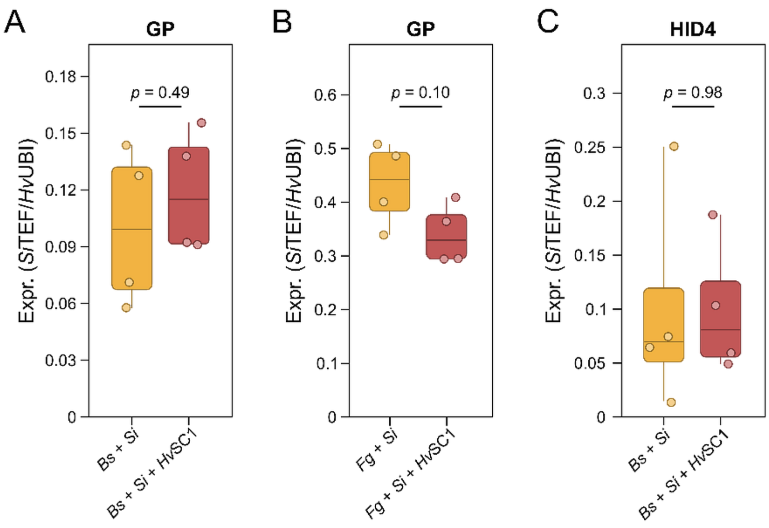


**Figure S10: Colonization of *Serendipita indica* in multi-kingdom contexts.** All plots show the relative expression of *S. indica* TEF normalized to barley Ubiquitin (UBI) expression as a proxy for fungal colonization. A) *S. indica* colonization in GP roots co-inoculated with the fungal pathogen *Bipolaris sorokiniana* (Bs), in the absence or presence of HvSC1. B) *S. indica* colonization in GP roots co-inoculated with the fungal pathogen *Fusarium graminearum* (Fg), in the absence or presence of HvSC1. C) *S. indica* colonization in HID4 roots co-inoculated as described in A. Statistical significance was assessed using a Kruskal–Wallis test followed by Dunn's multiple-comparison test ($p < 0.05$).
